# Heterogeneity and dynamics of ERK and Akt activation by G protein-coupled receptors depend on the activated heterotrimeric G proteins

**DOI:** 10.1101/2021.07.27.453948

**Authors:** Sergei Chavez-Abiega, Max L.B. Grönloh, T.W.J. Gadella, Frank J. Bruggeman, J. Goedhart

## Abstract

Kinases are fundamental regulators of cellular functions and play key roles in GPCR-mediated signaling pathways. Kinase activities are generally inferred from cell lysates, hiding the heterogeneity of the individual cellular responses to extracellular stimuli. Here, we study the dynamics and heterogeneity of ERK and Akt in genetically identical cells in response to activation of endogenously expressed GPCRs. We use kinase translocation reporters, high-content imaging, automated segmentation and clustering methods to assess cell-to-cell signaling heterogeneity. We observed ligand-concentration dependent response kinetics to histamine, a2-adrenergic, and S1P receptor stimulation that varied between cells. By using G protein inhibitors, we observed that Gq mediated the ERK and Akt responses to histamine. In contrast, Gi was necessary for ERK and Akt activation in response to α2-adrenergic receptor activation. ERK and Akt were also strongly activated by S1P, showing high heterogeneity at the single cell level, especially for ERK. In all cases, the cellular heterogeneity was not explained by distinct pre-stimulation levels or saturation of the measured response. Cluster analysis of time-series derived from 68,000 cells obtained under the different conditions revealed several distinct populations of cells that display similar response dynamics. The single-cell ERK responses to histamine and brimonidine showed remarkably similar dynamics, despite the activation of different heterotrimeric G proteins. In contrast, the ERK response dynamics to S1P showed high heterogeneity, which was reduced by the inhibition of Gi. To conclude, we have set up an imaging and analysis strategy that reveals substantial cell-to-cell heterogeneity in kinase activity driven by GPCRs.

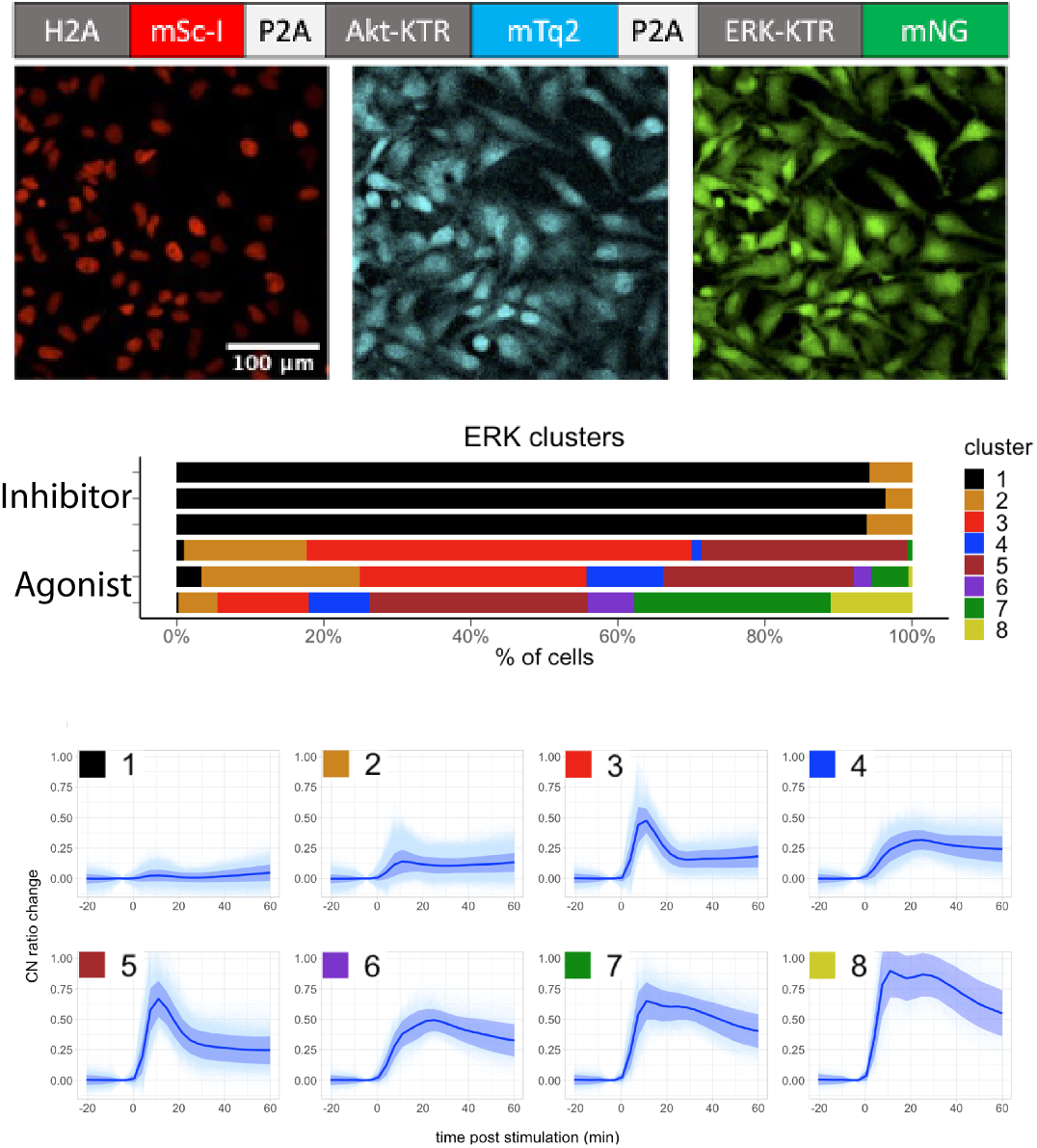

## INTRODUCTION

There are over 500 kinases encoded by the human genome, playing a fundamental role in regulating key biological processes within cells [1]. Kinases are major drug targets for oncology, with many approved drugs for the treatment of several breast and lung cancer types [2]. Kinases can either phosphorylate serine/threonine residues, or tyrosine, or in some cases both. The activity of kinases is regulated by events such as ligand-binding or phosphorylation by other kinases [3].

G protein-coupled receptor (GPCR) mediated signaling pathways involve many different kinases. The best characterized and studied kinases are PKA, PKC and Akt (or PKB) from the AGC family, and the mitogen-activated protein kinases (MAPKs) ERK, p38 and JNK. The activity of kinases such as PKA or PKC can be often tracked back to specific heterotrimeric G protein families. For instance, the relative activities of both Gas and Gai determine the cAMP levels in the cytosol [4], and cAMP modulates PKA activity by binding to its inhibitory domain [5]. Similarly, PKC is usually activated by increased levels of DAG and Ca^2+^, which occurs as result of PLCβ activation by Gq [6]. In contrast, the activity of kinases such as Akt or MAPKs is more downstream of the G protein-coupled receptor and, therefore, determined by different G proteins and pathways. The classic downstream effector of Gq is PKC, which can activate ERK. On the other hand, it is not so clear how Gq would affect Akt. The molecular network that connects the activity of Gi with kinases is also not so clear. In addition, other components involved in GPCR signaling, i.e. the β-arrestins that are traditionally considered exclusively as mediators of receptor internalization, can activate the MAPK ERK [7].

Traditionally, the kinase activities are inferred from cell lysates, hiding the heterogeneity of the individual cellular responses to extracellular stimuli. With the advent of genetically encoded biosensors, individual cells can be tracked over time. Several fluorescence-based biosensors are available that report kinase activity, each with different designs, fluorophores, read-outs, dynamic range, and sensitivity [8]. We decided to use kinase-translocation reporters (KTRs) because of the flexibility in the choice of the fluorophore and because they use a single channel [9]. KTRs suitable for monitoring of MAPKs and Akt have been described [10,11]. Since aberrant behavior of ERK and Akt is found across cancer types, these kinases are heavily investigated as potential therapeutic targets [12].

Single-cell studies on GPCR signaling pathways are still scarce, and the majority of studies on ERK activity in single cells are restricted to the study of growth factors. ERK activation kinetics is known to be very dynamic and to vary greatly between growth factors and concentrations [13]. Such studies indicate that cell subpopulations can be identified on the basis of single-cell responses. Therefore, we decided to (i) examine whether KTRs are sufficiently sensitive to detect activation of endogenous GPCRs and (ii) study the contribution of the Gq and Gi protein families to the activities of the kinases ERK and Akt in single cells.

## RESULTS

### Establishing a cell line for detection of ERK and Akt activation with translocation reporters

To investigate the relation between heterotrimeric G proteins and the activities of ERK and Akt in single cells, we employed kinase translocation reporters (KTRs). To detect Akt and ERK, we used Akt-FoxO3a-KTR [14] tagged with mTurquoise2 (mTq2) [15], and ERK-KTR [9] fused with mNeonGreen (mNG) [16]. To facilitate the identification of nuclei, we added a histone tagged mScarlet-I (mScI) [17]. The open reading frames of the three components were connected with P2A sequences, which ensures quantitative co-expression of the three proteins from a single open reading frame. The plasmid is named HSATEN (<u>H</u>istone-<u>S</u>carletI | <u>A</u>kt-KTR-m<u>T</u>urquoise2 | <u>E</u>RK-KTR-m<u>N</u>eonGreen). HeLa cells expressing HSATEN are shown in Figure 1.

**Figure 1.**
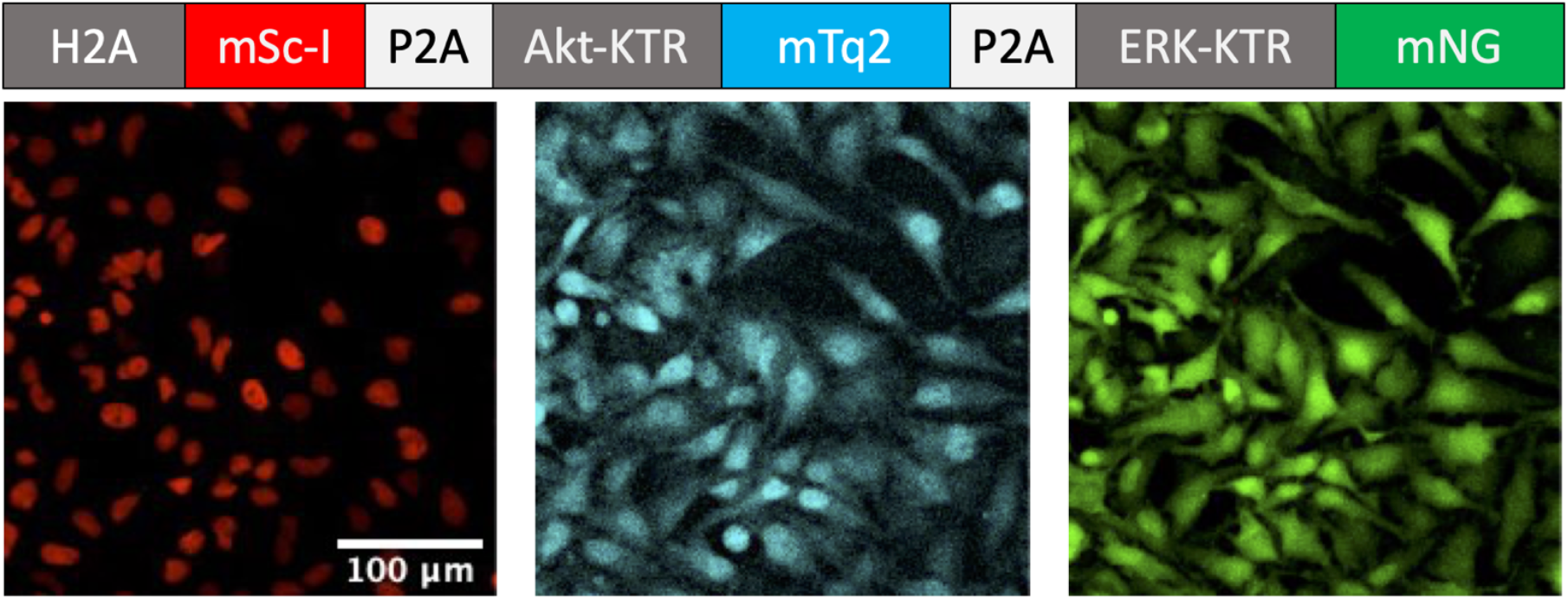
Triple color reporter plasmid. Top: A schematic drawing of the open reading frame of the construct with histone 2A (H2A) tagged with mScarlet-I (mSc-I), the Akt kinase translocation reporter (Akt-KTR) tagged with mTurquoise (mTq2) and the ERK kinase translocation reporter (ERK-KTR) tagged with mNeonGreen (mNG). The P2A sequences ensure separation of the proteins. Bottom: HeLa cells expressing the construct. From left to right: The nuclear marker, Akt-KTR, and ERK-KTR in red, cyan, and green respectively.

We used the PiggyBac transposon system [18] to generate cells that stably express the triple color reporter, and used fluorescence-activated cell sorting (FACS) to isolate single cells and obtain monoclonal populations. As can be seen in Supplemental Figure S1, ~60% of the cells were positive for mNeonGreen and mScarlet-I. Using the green fluorescence intensity levels, we sorted cells with intermediate levels of fluorescence in 4 pools. Next, we characterized the KTR response to fetal bovine serum (FBS), which strongly activates growth factor signaling and kinase activity.

To quantitatively compare the responses, we set up an analysis pipeline that quantifies the ratio of the cytoplasmic to nuclear intensity (C/N) of single cells, reflecting the kinase activity [14]. The pipeline uses FIJI [19] for background correction, CellProfiler [20] for segmentation, and the R programming language [21] for processing and visualizing the data. Supplemental Figure S2 shows the different steps of the analysis procedure. The scripts and fully reproducible instructions are available (https://github.com/JoachimGoedhart/Nuclear-translocation-analysis). This analysis pipeline is used for all data presented in the manuscript.

Based on the KTR responses, we decided to continue with pool 3. To examine whether the ERK and Akt basal levels could be reduce by serum starvation, we replaced the growth medium with serum free imaging medium and followed the C/N ratio over time. A reduction of the C/N ratio is observed and this reaches a plateau after ~100 minutes (Supplemental Figure S4). All of the following experiments were performed ~ 2hrs after replacing the medium to reduce the basal activity of ERK and Akt.

Next, we examined the effect of the MEK inhibitor PD 0325901. Pre-incubation with the inhibitor for 20 minutes blocked the response of the ERK-KTR to FBS, but not that of Akt-KTR (Supplemental Figure S5). This supports previous observations [14] [15] that the P2A effectively separates the different components, since the Akt-KTR and ERK-KTR show independent relocation patterns.

Pool 3 was used to isolate several clones. The monoclonal cell lines were tested for their response to FBS and the fluorescence intensity of the biosensors was quantified. The data is shown in Supplemental Figure S3 and the selection is discussed in Supplemental note 1. A single clone was selected and used for the remainder of our studies.

### Activation dynamics of ERK after GPCR activation

We selected three G protein-coupled receptor families, based on their capacity to activate different families of heterotrimeric G proteins and their expression in HeLa cells. We selected histamine receptors (HRs) [22,23] and sphingosine-1-phosphate receptors (S1PRs) [24] for which we used the respective endogenous ligands histamine and S1P. We also selected α2-adrenergic receptors (α2ARs) [25] and used UK 14304 (UK), also called brimonidine, a widely used full agonist with very high potency and selectivity [26]. To examine the activation of ERK by the three different GPCRs, we added a saturating concentration of agonist to the HeLa cell line expressing the KTR reporter. All three agonists were capable of inducing an increase in ERK activation as measured by an increased cytoplasmic/nuclear ratio. The responses were transient and showed considerable heterogeneity in amplitude (Figure 2).

**Figure 2.**
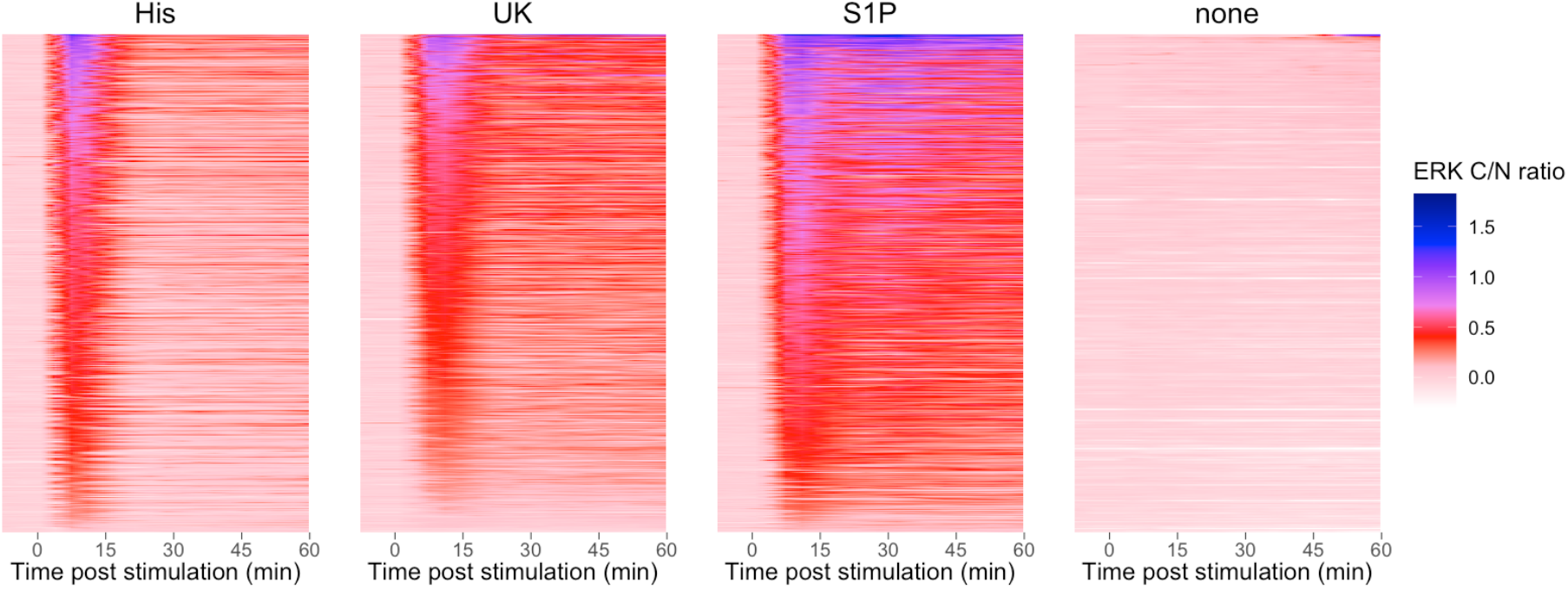
Single-cell time-lapse ERK responses to maximum ligand stimulatory concentrations. HeLa cells were treated with 100 μM Histamine (His), 100 pM UK 14304 (UK), 1300 nM sphingosine-1-phosphate (S1P) or no ligand (none). The ligand was added at t=0 and remained present. The ERK C/N ratio is presented as a false color and reflects the cytoplasmic over nuclear ratio of the ERK KTR, normalized by subtracting the average from the two time points prior to stimulation. For each ligand, the data corresponds to at least three biological replicates which are combined and sorted according to their integrated response.

### Concentration-response curves of ERK activation

Next, we examined the effect of different concentrations of agonists. Histamine stimulation causes a transient increase in ERK activity from concentrations as low as 0.19 μM, as shown in Supplemental Figure S6. The maximum activity is concentration-dependent and is reached about 10 min post stimulation. Similarly, UK addition led to a rapid increase in ERK activity and reaches a transient maximum 10-15 min post stimulation, after which it decreases to reach a plateau 30 min later (Supplemental Figure S7). The increase is observed with concentrations as low as 0.41 pM. In contrast to histamine and UK, S1P showed a more complex pattern with peaks at different time points, depending on the concentration of the agonist (Supplemental Figure S8). Overall, the ERK activity was concentration dependent for all three agonists with considerable heterogeneity at all of the tested concentrations.

We used the ERK-KTR data to fit concentration-response curves for ERK activity using the area under the curve (AUC) as the measure of the response, which we calculated as the sum of C/N ratios between 7- and 38.5-min post stimulation. For each biological replicate at every concentration, we calculated the average AUC, indicated by a large dot in Figure 3. The average of the biological replicates was used to fit the curve, and the results are shown in Figure 3 and Supplemental Table T2. The EC50 values for histamine, S1P and UK were respectively 0.3 μM, 64 nM and 2.5 pM.

**Figure 3.**
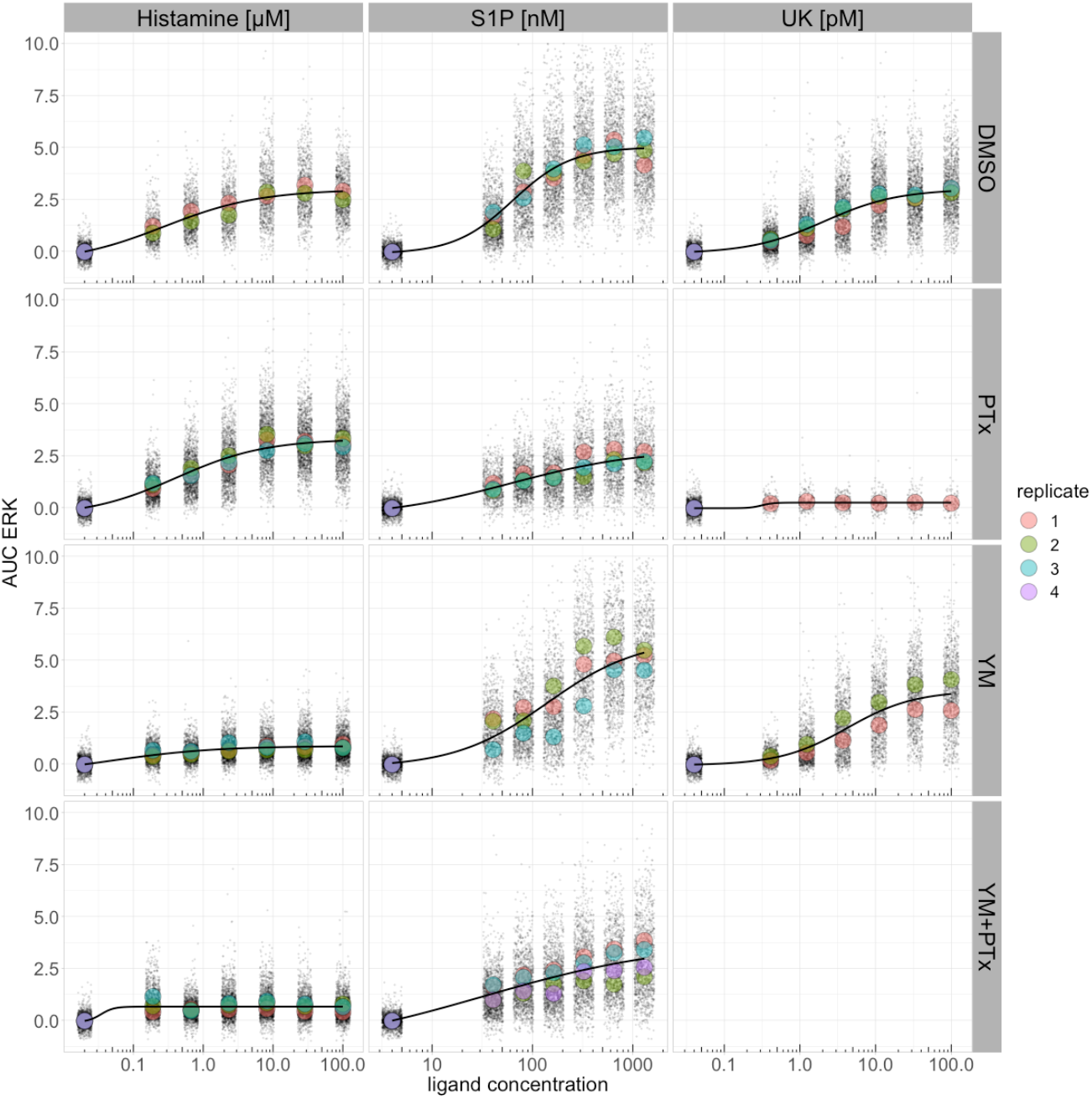
Concentration-response curves for ERK under all experimental conditions using the area under the curve (AUC) as the measure of response. The AUC was calculated as the sum of normalized C/N ratios from the time points 9-18, corresponding to 7 to 38.5 min post stimulation. The data were fitted with a four-parameter logistic equation, using the average of the average ERK AUC per biological replicate. Biological replicates are represented by different colors and their average is shown as a large dot.

### Effect of inhibiting heterotrimeric G proteins on ERK and Akt activation

To examine the role of heterotrimeric G proteins in the activation of ERK and Akt, we used YM-254890 (YM) to inhibit Gq and pertussis toxin (PTx) to inhibit Gi [27]. After inhibitor treatment, we stimulated the cells with histamine, S1P, or UK in a range of concentrations. The dynamics of the responses are reported in Supplemental Figures S6–S9. The AUC was used to construct concentration-response curves, and these are depicted in Figure 3. We note that YM, which targets Gq, inhibits the ERK response by histamine, whereas the response to UK is largely inhibited by PTx, which interferes with Gi signaling. The response to S1P is hardly affected by YM, but the amplitude is reduced by PTx.

Next, we examined the responses of Akt, which is simultaneously measured. The Akt responses were noisier due to lower amplitudes. Figure 4 shows the Akt responses to histamine. In the absence of inhibitors, the Akt activation is partially transient, with the response peaking 10 min post stimulation, and decreasing in the following 25 min to reach baseline levels (Figure 4A). Gi inhibition appears to cause a small increase in maximum activity and possibly a short delay in time of maximum activity, as shown in Figure 4B. Inhibition of Gq (figure 4C) decreases the maximum activity up to ~70%, and simultaneous inhibition of Gq and Gi causes a decrease of the responses up to ~90%, as shown in Figure 4D. These Akt amplitudes and effects of inhibitors are largely similar to those observed for ERK.

**Figure 4.**
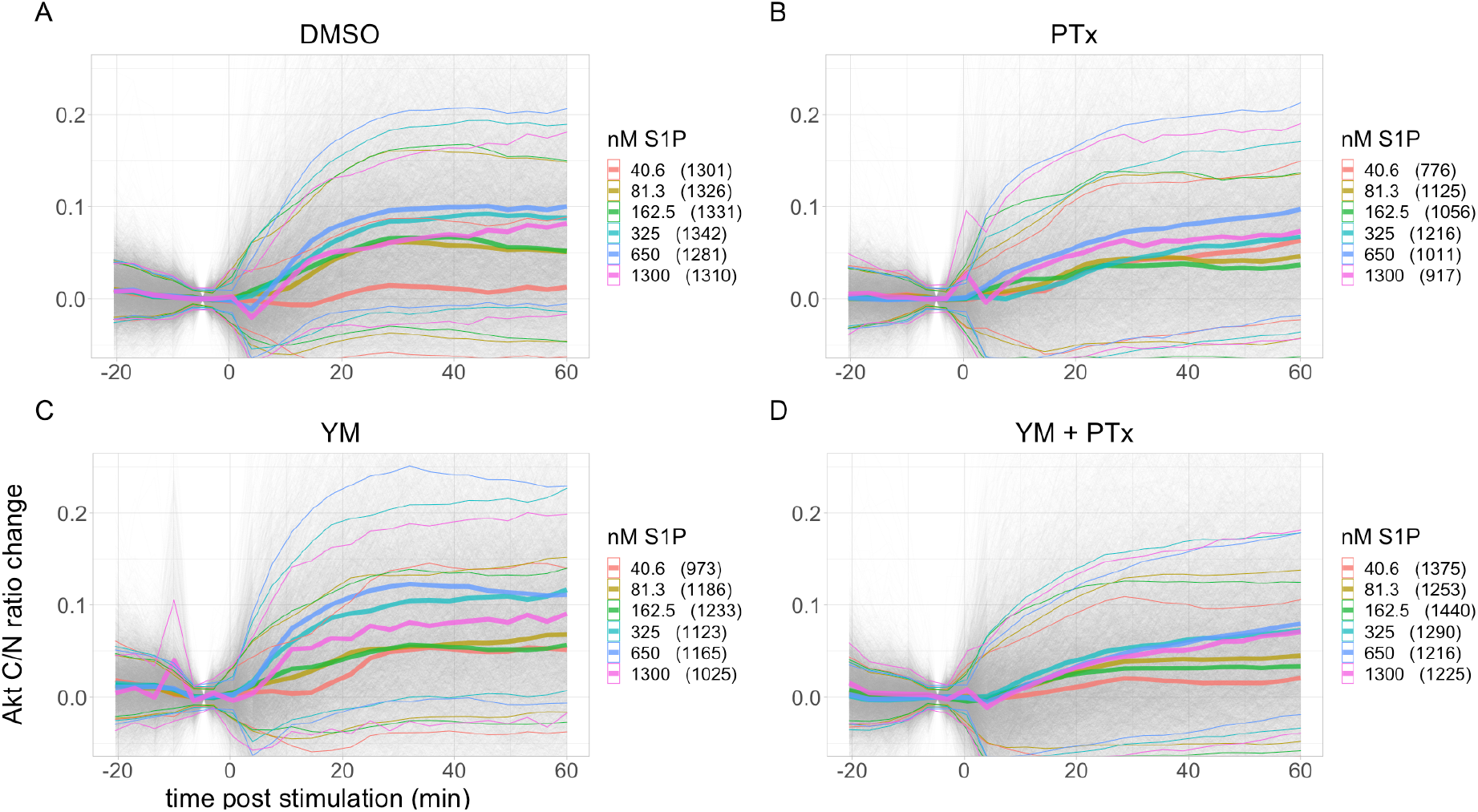
Akt responses to different concentrations of histamine and the effect of Gq and Gi inhibition. A: No inhibitor (DMSO). B: Gq inhibition (YM). C: Gi inhibition (PTx). D: Combined Gq and Gi inhibition (YM+PTx). Akt C/N ratio change is calculated by subtracting the average from the two time points prior to stimulation. Each panel shows combined data from at least three biological replicates. Gray lines represent single cell traces. Thick colored lines show the mean and thin colored lines the standard deviation for each ligand concentration. Number of cells are shown between brackets.

### ERK and Akt activities are correlated

It is striking that Gq inhibition has a similar inhibitory effect on Akt and ERK activity when cells are treated with histamine. To examine the correlation between ERK and Akt activity in more detail, we calculated the integrated response (AUC) for ERK and Akt in every cell for the different treatments. By plotting the ERK versus Akt activity, the relation between both activities can be visualized. As can be inferred from Figure 5, there is a moderate positive correlation between both kinase activities for each ligand. In conditions where G protein inhibition drastically affects the activity of the kinases, such as YM for histamine and PTx for S1P, the ERK responses are more strongly reduced than the Akt responses. Finally, we note that for S1P, the Akt activity is hardly or not reduced in the presence of inhibitors.

**Figure 5.**
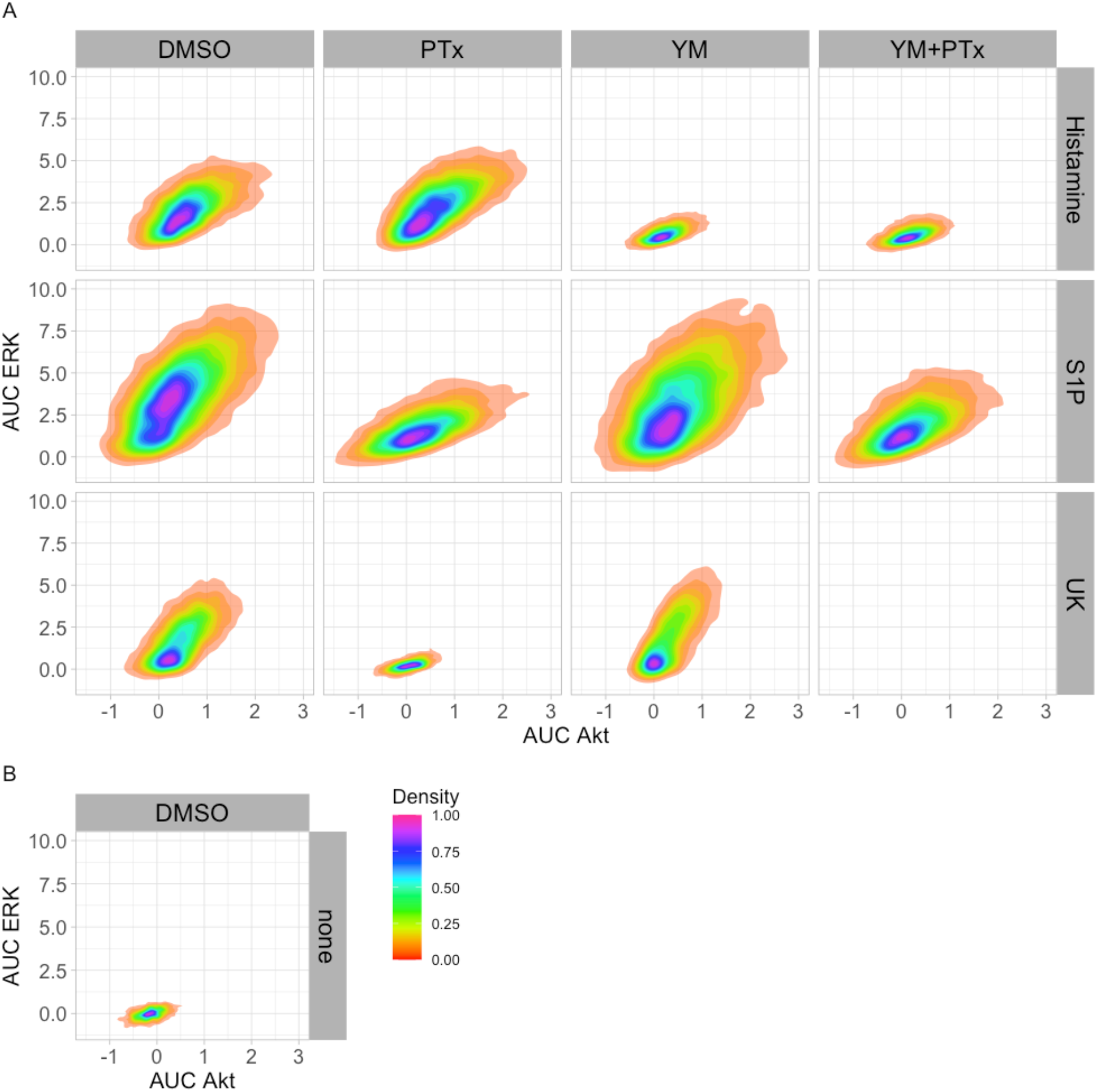
Activity of ERK versus activity of Akt per cell. AUC is used as the measure of response and was calculated as the sum of normalized C/N ratios from the time points 9-18, corresponding to 7 to 38.5 min post stimulation. Saturating concentrations of the ligands were used. For each cell, the AUC of Akt was plotted against the AUC of ERK, and the data from all biological replicates per condition is shown.

### Basal kinase activity does not affect the response amplitude

The measured single-cell kinase activities within an experimental condition, or even within a biological replicate, exhibit considerable heterogeneity. This can be clearly observed in the data shown in Figures 2–5. A possible explanation for the observed heterogeneity is that differences in basal kinase activity affect how the cells respond to the stimulus.

The information on basal kinase activity is lost when the data are normalized to set the initial C/N ratio to unity. To examine how the initial C/N ratio affects the response dynamics, we looked at the original, non-normalized data. Figure 6A, shows the variability among the C/N ratios for each KTR before ligand stimulation. For ERK, these start C/N ratios are spread evenly between 0.20 to 0.75, whereas for Akt most are within 0.30 and 0.65.

**Figure 6.**
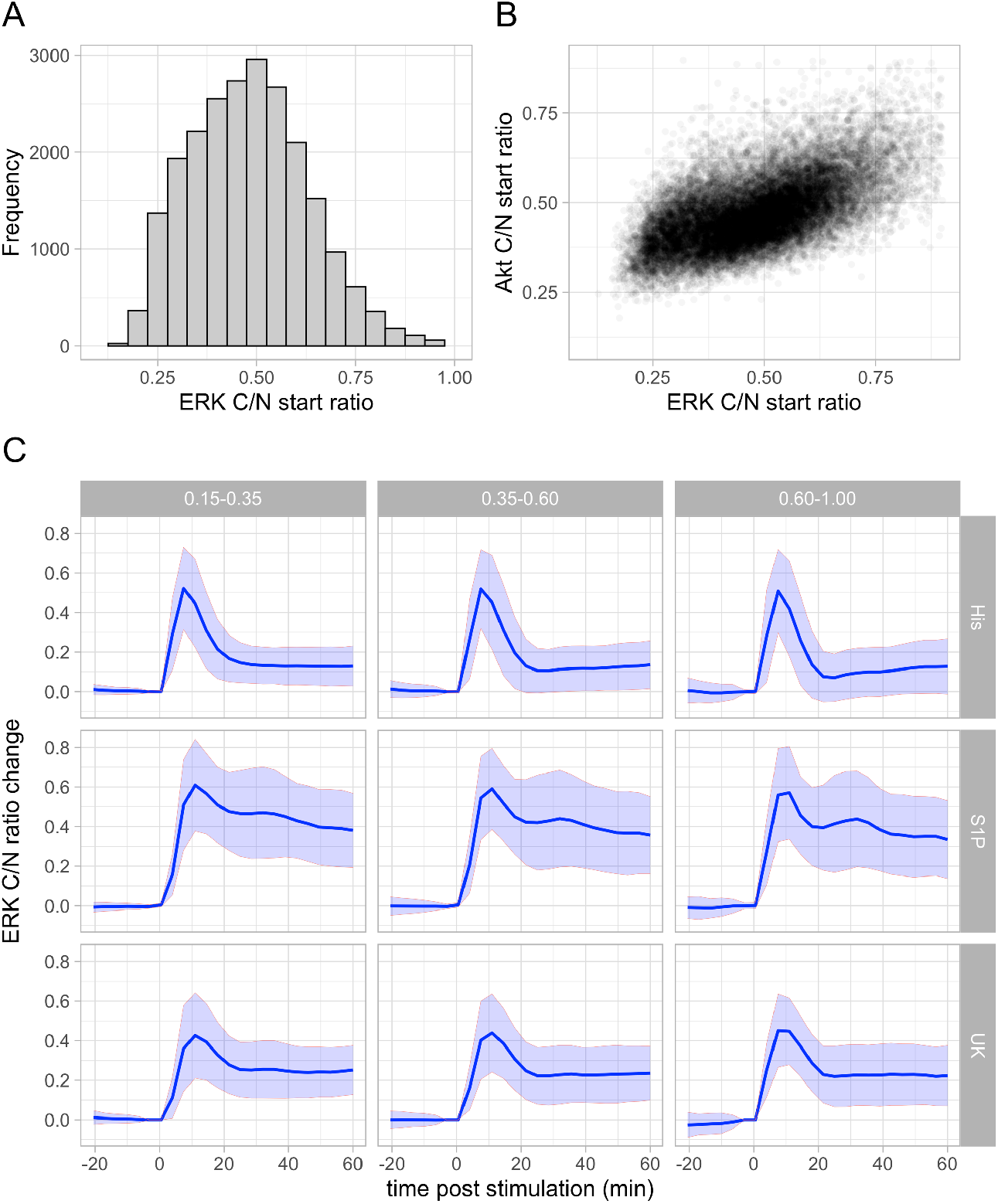
Distribution of C/N start ratios and effect on the ERK response. (A) Frequency of the average C/N ratios of ERK prior to ligand stimulation, using the data from single-cells from all the experiments. (B) Relation of the resting ERK C/N ratios and the resting Akt C/N ratios. (C) The data for the ERK responses at the maximum concentration of each of the ligands was grouped according to the start ratio, as indicated in the labels on top of the graphs. The ‘low’ pool had a start ratio 0.15-0.35, the middle pool a start ratio of 0.35-0.6 and the ‘high’ pool a start ratio of 0.6-1.0. The ERK C/N ratio change was normalized by subtracting the average of the two time points prior to stimulation. The line shows the average and the ribbon shows the standard deviation.

To examine whether the start C/N ratios, that reflect basal kinase activity, have an effect on the absolute C/N ratio changes, we decided to split each of the 3 datasets (without inhibitors) into 3 groups that represented relatively low, medium, and high start ratios. Figure 6B show the results for ERK and Supplemental Figure S10 for Akt. Overall, cells with different start ratios show comparable curve shapes and maximum activity, for the three ligands and for both kinases. For the lowest concentrations, there appears to be a trend where the ERK maximum activity increases slightly with the start ratio, but the differences are relatively small.

To conclude, our data show that the absolute changes in C/N ratios are hardly or not affected by the start C/N ratios. This suggests that the measured biosensor responses are not saturated in our experiments and that we can capture the entire range of kinase activities.

### Clustering reveals different kinase responses to GPCR activation

To get more insight in the heterogeneity and possible patterns in the response, we turned to cluster analysis. Clustering simplifies the data by defining different categories that group mathematically similar responses. This method had previously been used to examine the response of FRET biosensors [28] and KTRs [29]. First, we used a subset of the data to explore the optimal clustering method and to find the optimal number of clusters.

From our data, it is clear that there are major differences between the quantified ERK and Akt responses. First, the dynamic range of the ERK responses to the ligands is ~3-4 times bigger than that of Akt. Second, ERK displays various different curve shapes, whereas Akt responses vary almost exclusively in amplitude. Third, due to the low dynamic range, small variations in the focal plane during imaging can have a significant effect on the Akt ratios. For these reasons, we decided to evaluate 3-5 clusters for Akt, and 8-10 clusters for ERK. We consider that these cluster numbers capture most of the variability in the data, without complicating interpretation of the results, and providing with high-quality meaningful information. In addition, we chose to use the C/N ratio changes between 7- and 38.5-min post stimulation, as it contains most of the information.

Due to popularity for trajectory analysis and access to clustering programming packages, we chose to use k-means clustering and hierarchical clustering. After applying the different clustering methods to a subset of the combined data from all ligands and conditions, we used several metrics to assess and compare the quality of the clustering methods. The advantage of considering several metrics is that we reduce the risk of picking a cluster number that may be favored by a single indicator, but not by the rest. For each of the metrics, the higher the output, the better the quality of the result.

Supplemental Figure S11, shows the metrics for various cluster numbers for the ERK and Akt data respectively. As can be observed, the multiple metrics do not always show similar trends, which is not surprising given the differences in the ways they are calculated. In order to combine the different metrics to select clustering candidates, we decided to normalize each of the metric values by dividing it to the highest value among all the 15 combinations. Then, for each combination (cluster method and number) we added the values from all metrics, and the results are shown in Supplemental Table T3. From these, we picked two combinations per kinase, shown in blue, based on a higher score and lower number of clusters. Finally, we generated two plots to inspect the selected clustering approaches. First, the distribution of the clusters among the negative controls (no ligand added) and the experiments with the highest ligand concentrations. Second, we plotted the trajectories per cluster, using all 15 000 cells. These plots are shown in Supplemental Figures S12 and S13 for ERK and Akt respectively. Both algorithms yielded similar results and we decided to use 8 clusters for ERK with the Manhattan distance and Ward2 linkage method. Similarly, for Akt, we chose 3 clusters based on the Euclidean distance with Ward2 linkage.

Once we had selected the clustering method and optimal number of clusters, we applied it to the combined data from all ligands and inhibitory conditions (~68,000 cells). Figure 7 shows 8 distinct response patterns for ERK activation, including no/low responses (cluster 1 and 2), transient responses (cluster 3 and 5) and different patterns of a more sustained response (cluster 4, 6, 7 and 8).

**Figure 7.**
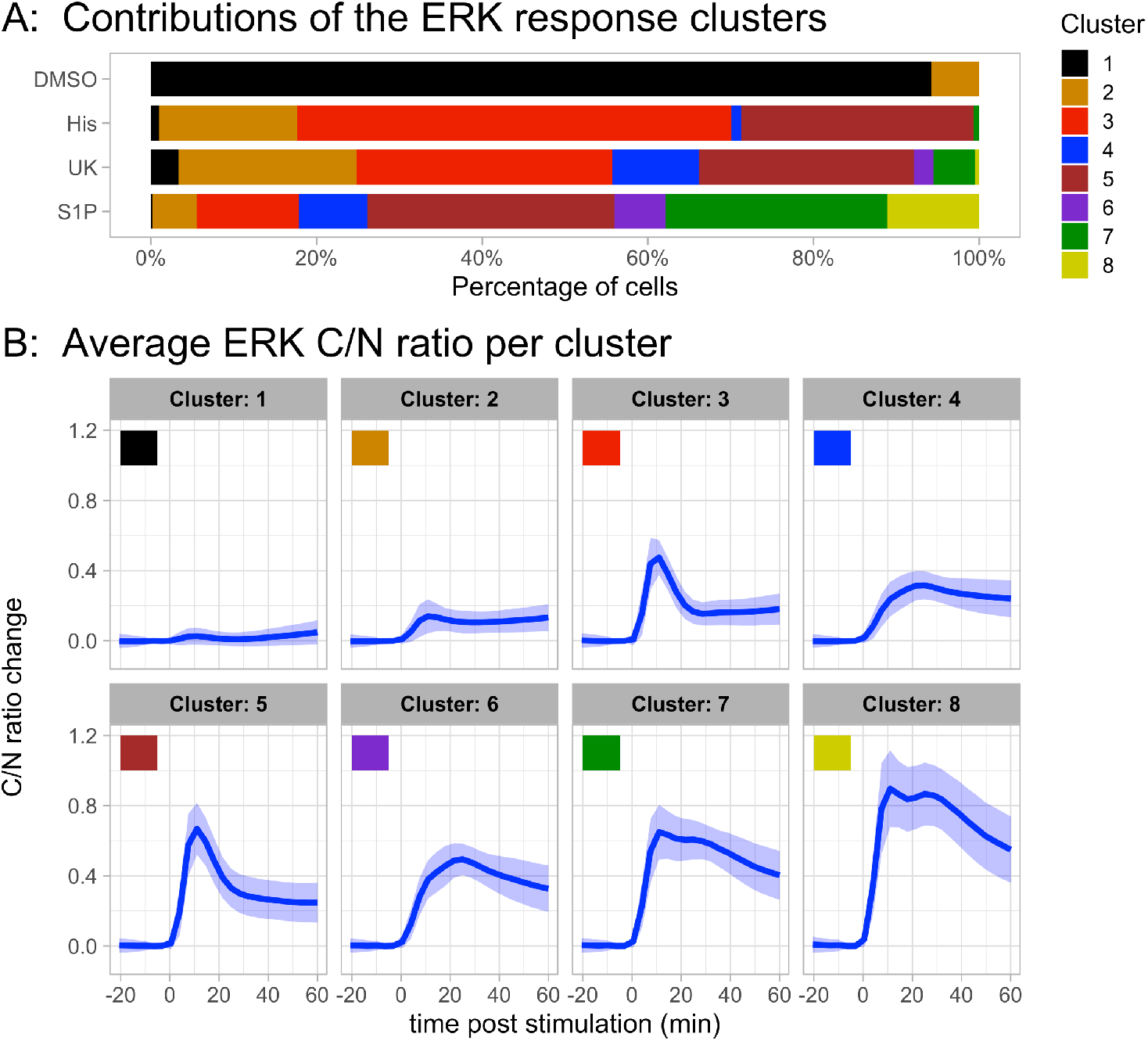
Results of clustering all data for the ERK responses. The selected method has 8 clusters and uses Manhattan distance and the Ward2 linkage method. It was applied to all the cells from the combined experiments with different ligands, concentrations, conditions, and negative controls. (A) Cluster distribution of responses in a control and in the condition of maximal ligand concentration. The control reflects addition medium instead of ligand. (B) Average trajectory and frequency of each cluster. Per cluster, the lines represent the average trajectory and the ribbon the standard deviation.

**Figure 8.**
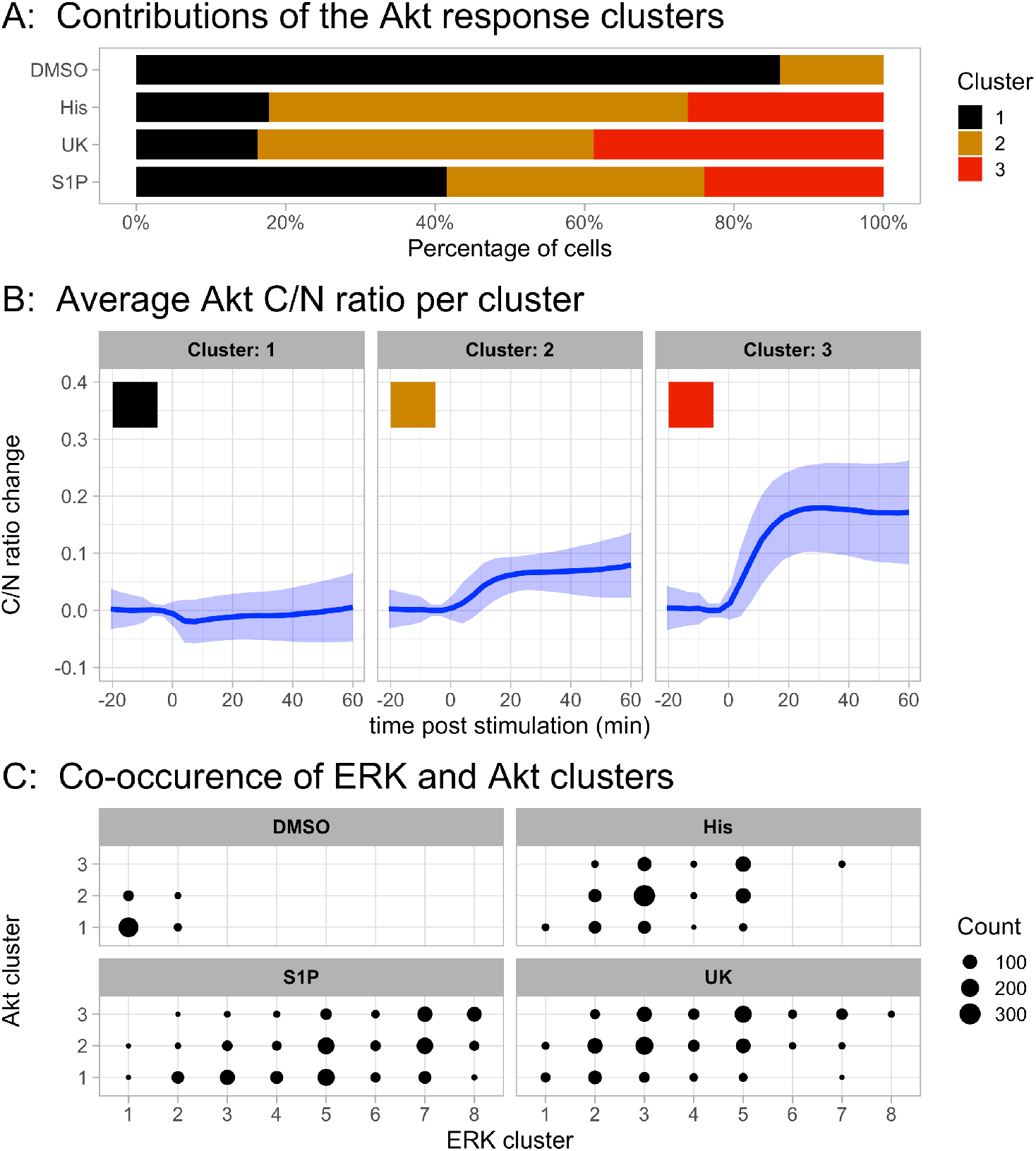
Results of clustering all data for the Akt responses. The selected method has 3 clusters and uses Euclidean distance and the Ward2 linkage method. It was applied to all the cells from the combined experiments with different ligands, concentrations, conditions, and negative controls. (A) Cluster distribution of responses in a control and in the condition of maximal ligand concentration. The control reflects addition medium instead of ligand. (B) Average trajectory and frequency of each cluster. Per cluster, the lines represent the average trajectory and the ribbon the standard deviation. (C) Co-occurrence of the ERK and Akt clusters for the control condition and for each of the three ligands at maximal concentration.

Our initial qualitative judgement that the response to histamine and UK is similar, is also quantitatively supported by the graph in Figure 7A that shows the contribution of each cluster to a treatment. A transient response dominates for these agonists. In contrast, the response to S1P is very heterogeneous, with contributions of cells that show transient ERK activity and cells that show sustained activity. The biphasic ERK activation pattern, which is specific for stimulation with S1P are reflected by clusters 7 and 8.

The cluster analysis for Akt is shown in Figure 8. The responses are grouped in three patterns, one of non-responding cells and two with sustained responses, differing in amplitude. The activation of Akt is remarkably similar between the different treatments. In figure 8C, the co-occurrence of ERK and Akt clusters is depicted. Also in this plot, there is similarity between the responses to the ligands histamine and UK, with a high cooccurrence of transient ERK activation (cluster 3 & 5) with a sustained Akt response (cluster 2&3). The response to S1P shows again a larger heterogeneity.

Since the ERK activation shows the largest heterogeneity, we examined the effect of heterotrimeric G protein inhibition. For each of the conditions, we display the relative contribution of each of the different patterns. The results are depicted in Figure 9 and Supplemental Figure S14. The results for Akt are shown in Supplemental Figures S15–S16.

**Figure 9.**
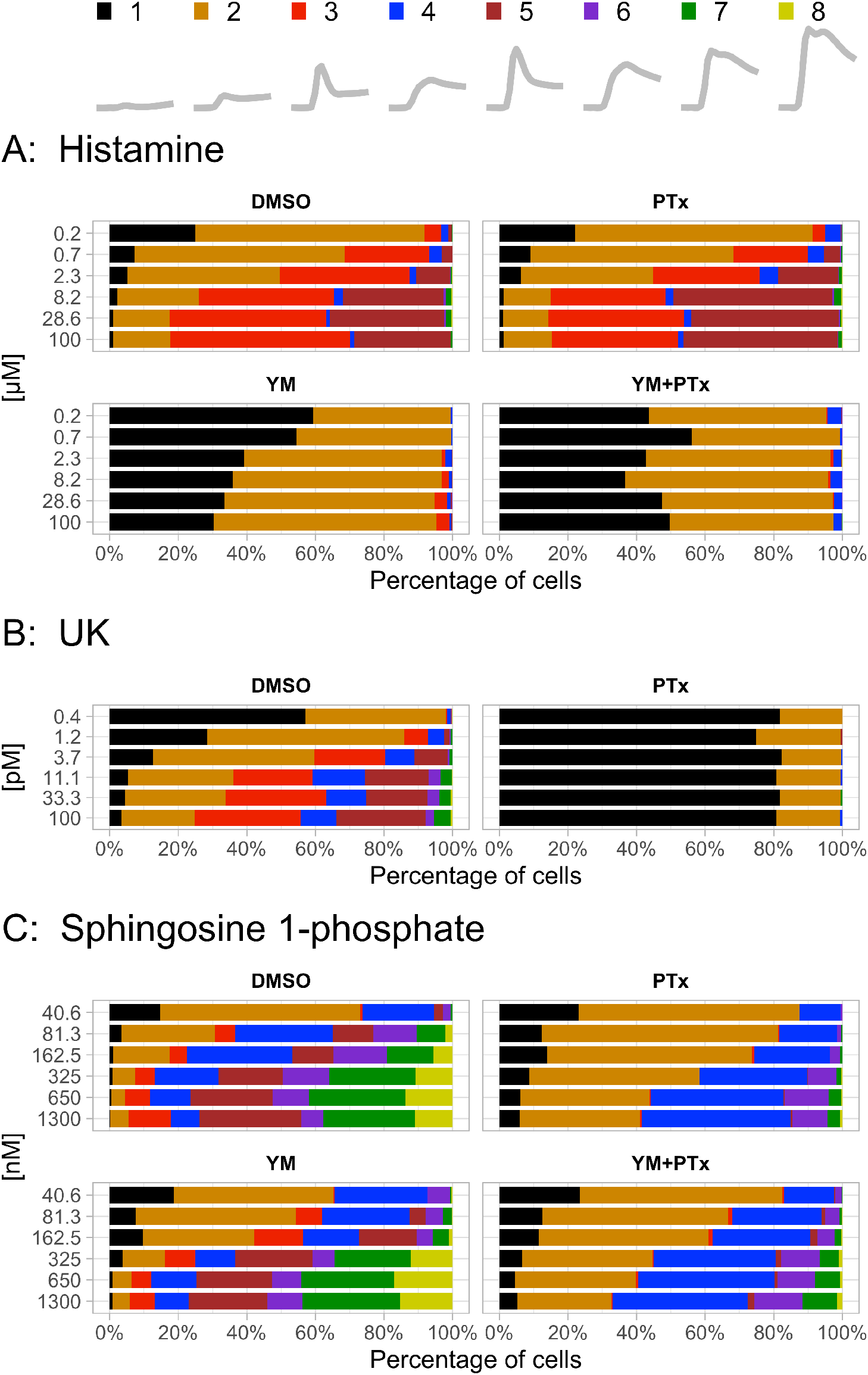
Cluster distribution of ERK responses at different concentrations per ligand. (A) The temporal profile of each cluster (same as figure 7B) and the corresponding color code. (B) Histamine, (C) UK 14304 and (D) Sphingosine 1-Phosphate. For each ligand, the panels represent the different experimental conditions. No inhibitor (DMSO), Gq inhibition (YM), Gi inhibition (PTx), and combined Gq and Gi inhibition (YM+PTx).

The fraction of non-responding cells (cluster 1) systematically decreases when the ligand concentrations of histamine, UK or S1P are increased. Although the inhibitors are effective in some combinations, i.e. YM and histamine, PTx and UK, there is still a fraction of cells that respond. This suggests that the ERK activity is not exclusively due to the activation of the corresponding heterotrimeric G protein.

We note that there are hardly any unresponsive cells for S1P concentrations above 81.3 nM. However, there is substantial heterogeneity in the responses above this concentration. At least 6 different response patterns can be discerned. Surprisingly, the heterogeneity is strongly reduced when Gi signaling is inhibited by PTx. The rapid rise in ERK activation (as observed for stimulation by histamine and UK) is abolished and a delayed and more sustained response is the result. The effect of YM on the ERK activity is weak. To summarize, cluster analysis reveals the contribution of different ERK activation patterns and the palette of patterns can be profoundly changed by inhibition of heterotrimeric G proteins.

## DISCUSSION

Most studies of kinases activated downstream of GPCR signaling pathways are performed using biochemical assays on cell populations. These methods cannot measure the dynamics in individual cells and detect the heterogeneity of the individual responses. The recent engineering of fluorescent biosensors that are based on translocation has enabled high-content imaging of kinases such as ERK, Akt, JNK and p38. These reporters have been successfully used to study growth factor signaling in a number of settings and systems [30,31]. So far, only a couple of studies looked into kinase activation by GPCRs in single cells with KTRs and these studies used overexpressed receptors [32,33].

Here, we use KTRs that report on ERK and Akt [14] to generate monoclonal stable cell lines that can be used for multiplex imaging and demonstrate that the KTRs are sensitive enough to detect activation of endogenous GPCRs. This is in marked contrast to other fluorescent biosensors that, in our hands, typically require an overexpressed receptor for robust responses [34]. Our imaging pipeline enables high-content imaging of the responses, yielding quantitative, dynamic data from thousands of cells. The data was used to generate concentration-response curves for three agonists from the imaging data and to examine the effect of G protein inhibition. The analysis revealed different dynamics between GPCRs and the cluster analysis showed differences between subpopulations of cells activated with the same agonist.

Our initial idea was to use KTRs as specific read-outs for heterotrimeric G protein activity, which is relevant for understanding ligand-biased activation [35]. This would be achieved when Gq activation is linked to ERK and Gi activation results in Akt activity. We selected three agonists that would activate three different families of GPCRs that are endogenously present in HeLa cells. Histamine is reported to predominantly activate Gq in HeLa cells by the histamine H1 receptor [36] and UK activates Gi by α2-adrenergic receptors [37]. Our data with the inhibitors YM and PTx, which are selective of Gq and Gi respectively, show that the two agonists indeed preferentially activate a single heterotrimeric G protein class. Despite the activation of different heterotrimeric G protein families, the responses of the ERK-KTR to histamine and UK are remarkably similar. Both agonists also show a comparable effect on the amplitude and kinetics of the Akt-KTR. Therefore, our choice of KTRs does not enable the discrimination of signaling through Gi and Gq. The combination of an ERK- and Akt-KTR is not optimal, since their activities are largely correlated and similar for different G protein classes. Therefore, the measurement of Akt does not add information. Moreover, the Akt response had a relatively poor amplitude.

The situation for S1P is different. S1P can activate a number of different GPCRs, all known to be expressed by HeLa cells as shown in the supplemental figure S4A of [24]. As a consequence, S1P will activate a number of different heterotrimeric G protein families. We observed that activation of endogenous S1P receptors resulted in a strong, but highly heterogeneous ERK-KTR response, with two peaks in a population of cells. Both the dynamics and the amplitude varied between populations of cells and cluster analysis was applied to define 8 different patterns (including a flat line for non-responding cells). At least 6 of these patterns were identified at the higher S1P concentrations. From these data it is clear that genetically identical cells can respond in a highly heterogeneous manner to a single ligand, which is in line with previous studies [38]. Intriguingly, the heterogeneity in ERK dynamics is reduced when Gi signaling is inhibited. When PTx is present, the biphasic response is abolished and the first peak of activation is reduced, suggesting that the initial response is due to Gi signaling. This result demonstrates that by modulating the palette of heterotrimeric G proteins, the response dynamics are altered, which can be readily identified by cluster analysis. The clustering is a powerful method for the detection of patterns and simplification of large amounts of data. Yet, it should be realized that clustering is mathematical procedure that is not necessarily reflecting the biological processes. One example is the graded response of ERK and Akt activities to ligands, whereas cells are grouped in weak, middle and strong responders. This may be solved by developing and using clustering methods that take the underlying biological processes into account.

To enable a better insight in the specific heterotrimeric families that are activated by GPCRs, future studies looking into the response of different KTRs to different heterotrimeric G proteins and agonists are required. There is a translocation reporter, PKA-KTR, which is expected to be specific for Gs [9] and there are several KTR for which selectivity remains to be examined, e.g. p38-KTR and JNK-KTR. In addition, existing proteins that translocate in response to cell stimulation, including MRTF-A, YAP, NF-κB, and SMAD can be examined. The origins of the observed heterogeneity are unclear at present. We have used a monoclonal cell population and, therefore, the origin of the heterogeneity is likely non-genetic. In addition, we have verified that the differences between cells are not due to saturation of the sensors. Despite the use of monoclonal populations, gene expression is a stochastic process [39] and cellular noise resulting in differences in the relative concentrations of the components involved in the signaling network may lead to the observed differences [38]. Moreover, it is unclear what the consequences of this heterogeneity in kinase activities as a result of GPCR activation are. Heterogeneous single-cell responses dynamics has been previously linked to differences in physiologically relevant processes such as proliferation [40], metabolic adaptations [41], migration [42] and cell fate [43]. Since GPCRs are expressed ubiquitously and participate in many different processes, the implications of this heterogeneity need to be studied in a specific physiological context. Importantly, given the long-term cellular effects of ERK and Akt kinase activities, special attention should be given to changes in gene expression or cell cycle.

A limitation of our work is that the contribution of G protein independent mechanisms for ERK and Akt activation are unknown. At least two ways of activating ERK have been reported that may not require G proteins, i.e. β-arrestin mediated signaling [44] and transactivation of a receptor tyrosine kinases by a GPCR leading to ERK and Akt phosphorylation [45]. Based on our data, we cannot exclude that beta-arrestin or RTKs play a role in the activation of ERK and Akt. To study the role of non-classical routes to ERK activation, inhibitor studies, or probes that interrogate these processes would be useful. Increasing the number of probes to measure several processes simultaneously would provide a better picture of the contribution of different networks and their interactions. Multiplex, live-cell imaging with 6 probes has been demonstrated [46] and would enable the measurement of a reference for segmentation and 5 KTRs or other probes. Ongoing efforts to engineer brighter fluorescent proteins and hybrid genetic tags (e.g. HaloTags and SNAP tags) are important to further improve multiplex imaging. The functional translocation read-outs can potentially be combined with morphological profiling [47] for multiparameter, high-content imaging-based drug screens.

We hope that the new imaging strategy and analysis presented here, will be valuable for future studies that use imaging of kinase activity in single cells to connect GPCR activation with physiological effects.

## METHODS

### Reagents

S1P (Sigma-Aldrich, cat# S9666) was prepared as a 1.3 mM stock solution in methanol. Histamine (Sigma-Aldrich cat# H7125) was prepared as a 100 mM stock solution in water. UK 14,304 (Sigma-Aldrich cat# U104) was prepared as a 10 μM stock solution in DMSO. YM-254890 (FUJIFILM Wako Pure Chemical Corporation, cat# 257-00631) was prepared as a 1 μM solution in 33% DMSO in MQ water. Pertussis toxin (Invitrogen, cat# PHZ1174) was prepared as a 100 ng/mL solution in water. PD 0325901 (Sigma-Aldrich, cat# PZ0162) was prepared as a 1 mM solution in DMSO.

### Cloning single kinase translocation reporters and nuclear marker

The first step to generate the multicolor constructs was to clone the individual KTRs and the nuclear marker, and tag them with the FPs of interest. The ERK-KTR, developed by Regot et al. [9], and the Akt-KTR, by Maryu et al. [14], were part of the pHGEA plasmid kindly shared by Dr. Kazuhiko Aoki. Both sequences contained a P2A sequence in front, which we kept to ensure equimolar expression of the separate proteins from a single transcript [48].

P2A-ERK-KTR and P2A-Akt-KTR sequences were amplified by PCR from pHGEA (Akt: Fw 5’-TATA<u>GGTACC</u>AAACCATGGGGTCAGGGGCCACCAACTTC-3’ and Rv 5’-TATA<u>ACCGG</u>TATGCGGCCGCCGAGCGTGATGTTATC-3’, ERK: Fw 5’-TATAG<u>GTACC</u>AAACCATGGGGAGCGGGGCTACCAACTTC-3’ and Rv 5’-ATAT<u>ACCGGT</u>ATGCCGCCGGACGGGAATTG-3’) to introduce Acc65I and AgeI restriction sites (underlined in primers sequences). Next, the P2A-KTRs PCR products and Clontech N1 vectors containing either mTurquoise2 (mTq2) [15], mNeonGreen (mNG) [16], or mScarletI (mScI), [17] were digested with Acc65I and AgeI, ligated using T4 DNA ligase, and transformed by heat-shock using DH5α Escherichia coli competent cells.

In addition, the residues S294 and S344 in the Akt-KTR were mutated to Ala by site-directed mutagenesis (S294A: Fw 5’-CCAAGTGGCCTGGC<u>GC</u>CCCCACGTCACGCA-3’ and Rv 5’-TGCGTGACGTGGGG<u>GC</u>GCCAGGCCACTTGG-3’, S344A: Fw 5’-TGCGCCTCT<u>C</u>GCGCCCATGCTCTACAGCAG-3’ and Rv 5’-AGCATGGGCG<u>C</u>GAGAGGCGCATCATCGTCC-3’) as it has been reported that these residues in FOXO3 could be phosphorylated by ERK [49]. We used PfuTurbo DNA polymerase, followed by DpnI digestion to destroy template DNA.

To generate the nuclear marker, we replaced mTq2 from a Clontech N1 H2A-mTq2 for mScI using AgeI and BsrGI.

### Combining kinase translocation reporters

The second step was to combine the translocation reporters and the nuclear marker. Taking advantage of the compatible cohesive ends generated by digestion of Acc65I and BsrGI, we first generated P2A-Akt-KTR-mTq2-P2A-ERK-KTR-mNG by ligating P2A-ERK-KTR-mNG digested with Acc65I and P2A-Akt-KTR-mTq2 with Acc65I and BsrGI. Later, with the same approach we generated H2A-mScI-P2A-Akt-KTR-mTq2-P2A-ERK-KTR-mNG, which we will refer to as HSATEN. The plasmid is available from addgene (plasmid #129631).

To incorporate HSATEN into the piggyBac transposon vector pMP-PB [18], kindly shared by Jakobus van Unen and David Hacker, we digested both constructs with NheI and XbaI and then ligated them. Since the cohesive ends generated by these enzymes are compatible, we performed a colony PCR to determine which colonies expressed the construct in the right orientation. We used transposon vectors containing antibiotic resistance for puromycin, blasticidin, hygromycin, and zeocin. The plasmid with puromycin resistance was used in this study and is available from addgene (plasmid #129632).

### Cell culture

HeLa cells (American Tissue Culture Collection: Manassas, VA, USA) and HeLa stable cell lines were maintained in “full growth medium”, or Dulbecco’s Modified Eagle Medium with GlutaMAX (Gibco, cat# 61965059) supplemented with 10% fetal bovine serum (FBS) (Gibco, cat# 10270106), at 37°C in 7% CO_2_ in humidifying conditions. Cells were passaged every 2-3 days by washing with HBSS (Gibco, cat#14170), trypsinizing using 0.25% Trypsin-EDTA (Gibco, cat# 25200056), spinning down at 300 g for 5 min, and resuspending in full growth medium. All cells were routinely tested for mycoplasma by PCR.

### Generation of HSATEN cell lines

200 000 HeLa cells in 2 mL full growth medium were plated per well on a 6-well plate (Corning, cat# 3516) and let them grow overnight. The following day, we co-transfected 500 ng of pPuro-PiggyBac-HSATEN and 200 ng transposase using 3.5 μL PEI (1 mg/mL in water). As a negative control, we transfected HSATEN and transposase. 24h posttransfection, 1 μg/mL Puromycin (Gibco, cat# A1113803) was added to the cells, and after 48h, both media and puromycin were refreshed. After 72h of selection with puromycin, the cells were trypsinized and passed to T25 flasks until confluency.

To sort by fluorescence-activated cell sorting (FACS), the cells were first washed, trypsinized, spun down, and resuspended in full growth medium as for passaging. Then, the cells were spun down, resuspended in 2% FBS in HBSS containing 1 μg/mL DAPI (Invitrogen, cat# D1306), spun down, resuspended in DAPI-free 2% FBS-HBSS, and kept in the dark on ice. Cells were sorted with the FACSAria™ III (BD Biosciences, Franklin Lakes, NJ, USA), using a 100 μm nozzle at 20 psi pressure.

Single cells were identified by drawing gates using the area, width, and height of forward scatter (FSC) and side scatter (SSC), and living cells based on being DAPI-negative. Living cells were identified based on the DAPI staining. To draw the gates for mNG- and mScI-positive cells we used HeLa cells as a negative control. DAPI was excited with 405 nm and measured with a 450/50 nm bandpass emission filter. mNG and mScI were excited with 488 nm and 561 nm respectively, and detected with a 530/30 and 610/20 bandpass emission filters.

We selected 4 gates based on mNG intensity, distributed along the 50% brightest cells. We then sorted the pools into 15mL tubes, and single cells in 96-well plates. The tubes and plates contained full growth medium, supplemented with 10 mM HEPES and 1% Penicillin/Streptomycin (P/S) (Gibco, cat# 15140148). Additionally, the 96-well plates were first coated with 14 μg/ml fibronectin in PBS for 1h. The cells in the tubes were spun down, resuspended in full growth medium with 1% P/S, and seeded in wells or flasks, depending on the number of cells. The media of the 96-well plates was replaced the following day for full growth medium. The single clone populations were sequentially transferred to bigger wells/flasks to expand.

### Live-cell imaging

For live-cell imaging, we used the Leica TCS SP8 confocal microscope (Leica Microsystems, Wetzlar, Germany) equipped with a 10X air objective Plan Apo 0.40 NA and a Mercury lamp at 37°C. We excited mTq2, mNG and mScI with a 440 nm DPSS, 488 nm Argon, and 561 nm DPSS lasers. Fluorescence was detected using HyD detectors for mTq2 and mScI (452-500 nm and 590-675 nm) and a PMT detector for mNG (506-560 nm). The width of the detectors was controlled with sliders through the Leica Application Suite X (LAS X, Leica Microsystems, Wetzlar, Germany).

The day before imaging, ~120 000 cells were seeded in a glass bottom 8-well μ-slide (Ibidi, cat# 80827) in full growth medium. 2 h before imaging, the medium was removed and replaced with microscopy medium (MM) (20mM HEPES pH=7.4, 137mM NaCl, 5.4mM KCl, 1.8mM CaCl2, 0.8mM MgCl2, 20mM Glucose) containing 0.033% DMSO or 1 μM YM-254890 (dissolved in 33% DMSO in water). PTx was added at the end of the day the cells were seeded, at a concentration of 100 ng/mL. We incubated the cells for 2 h with serum-free MM prior to imaging to reduce the basal kinase activities of ERK and Akt.

The acquired images had a 12-bit color depth and 1024×1024 pixels resolution. Images were acquired every 3.5 min, and each image was the average of 4 frames.

To keep the cells in focus, we executed Best Focus on the first well at the beginning of each time-point, and the correction was extended to the rest of the wells. Ligands were pipetted to all wells during time point 7.

Ligand solutions were prepared in pre-warmed microscopy medium containing either DMSO or YM, depending on the experiment. 100 uL of each ligand solution was added to the well containing 300 uL. For histamine and UK, the stock solutions were used to prepare the solution with the highest concentration, and serial dilutions were prepared from it. For S1P, the serial dilutions were prepared in methanol using gastight syringes (Hamilton, #1702 and 1710), and medium was added afterwards up to 100 μL. Ligand solutions were kept at 37°C for 10 min before pipetting to avoid cellular stress. For stimulation with 5% FBS, 20 uL prewarmed FBS was added to each well containing 380 uL of MM.

To determine the concentrations that yield minimum and maximum Akt/ERK activities for each ligand, we tested concentrations in the following ranges: 0.13 - 200 μM, for histamine; 16 - 2600 nM, for S1P; 0.13 pM - 10000 pM, for UK.

### Image processing

A reproducible image processing pipeline using Fiji, CellProfiler and R is available as a GitHub repository: https://github.com/JoachimGoedhart/Nuclear-translocation-analysis. The repository includes example data, code, a manual and the expected outcome as a graph. Below, we describe the steps in detail.

Processing of the raw images was performed using FIJI [19]. The individual signals were not unmixed because the cross excitation and bleed through were close to zero. To facilitate segmentation of the nuclei, we first subtracted 250 counts from the mScI images to remove any counts in the cytoplasm due to overexpression of the H2A-marker. To remove the background from the mTq2 channel images, we applied a rolling ball of 70 pixels radius and used these images for quantification of the mTq2 signals. For identification of the cell boundaries, we first applied a Gaussian blur with sigma 2 to smoothen the mNG images and then applied a manual threshold from 300 to 65535 to obtain a binary mask.

### Segmentation and tracking of nuclei and cytoplasms

We used a custom-made CellProfiler (version 3.0.0) [20] pipeline for segmentation, measurement of intensity and shape features, and tracking. We first used the processed mScI images to identify the primary objects, i.e. nuclei. We used a global threshold of 330 counts to separate pixels into background and foreground, and included objects with a diameter within 8-20 pixels. Clumped objects were identified and separated according to intensity. To identify the cells, we used the nuclear regions of interest (ROIs) as seeds in the binary mNG images. The nuclear ROIs were expanded up to 5 pixels in all directions as long as there was no background. The cytoplasmic ROIs were simply determined as a subtraction of the nuclear ROI from the cellular ROI. The nuclear and cellular ROIs were then tracked through the time lapses. These ROIs were identified as unique objects if the distance between their positions in consecutive images was lower or equal to 3 pixels. Finally, the size/shape features of the nuclear and cytoplasmic ROIs were exported, together with the intensity features of these ROIs in the processed mTq2 and raw mNG images.

### Data processing

We then used a custom-made R [21] script to process the exported data from CellProfiler. First, we applied filters to exclude ROIs with mean intensity values lower than ~260 counts and higher than ~4000. In addition, we removed ROIs with an area lower than 100 pixels and average pixel radius of 1 or less. Then, we removed the objects that were not present in each time point as a single object. Finally, we calculated the cytoplasmic to nuclear intensity (C/N) ratio per cell by dividing the mean intensities of both ROIs, for mTq2 and mNG channels.

Due to the experimental setup, the imaging of the 6 wells is not simultaneous, as there is a delay of 0.5 min between each well and the subsequent. To get C/N ratios at the exact same times and simplify later analysis, we applied a linear interpolation to the data. In addition, data was normalized by subtracting the average of two time points prior to stimulation (usually the 5^th^ and 6^th^ time point) from every data point.

Area under the curve (AUC) was defined as the sum of the C/N ratios from the time points 9-18, corresponding to 7 to 38.5 min post stimulation.

### Concentration-response curves fitting

To estimate the EC50 for each condition, we fitted the data using a four-parameter logistic curve, with the function “drm” from the package “drc” [50]. For each concentration, we used the average of the average value per biological replicate. The response from the negative control was entered as a low concentration, as the log of 0 is undefined. The data and the R-script for fitting the data is available: https://github.com/JoachimGoedhart/GPCR-KTR

### Trajectories clustering with R

To cluster the data, we decided to combine the data from the three ligands, in order to compare the heterogeneity of responses among the three ligands. In addition, we included data from 3 experiments where only vehicle was added, to use as negative control. To speed up the analysis, we used a subset of 15 000 cells, equivalent to ~ 20% of the total number of cells. We used two different clustering approaches, hierarchical clustering, and k-means clustering, and applied these to the normalized ratios from the time points 9-18, corresponding to 7 to 38.5 min post stimulation.

For the hierarchical clustering, we first used the function “parDist”, from the package “parallelDist” [51], to create a matrix with the calculated “distances” between all the cells. These distances represent the (dis)similarity between any two trajectories, and we used two of the most commonly used distance metrics, Manhattan, and Euclidean. We then used the function “hclust” from the package “fastclust” [52] to cluster the trajectories according to the values in the distance matrix, using the linkage methods Ward and Ward2. The result is a dendrogram that can be cut into a k number of clusters or at a certain “height”. The Ward method is commonly used with squared Euclidean distances, but it can be used with non-squared Euclidean distances [53] or Manhattan distances [54]. The only difference between Ward and Ward2, is that Ward2 first squares all the given distances.

For k-means clustering, it is necessary to first indicate the number of clusters (k) to be used. The Euclidean distances are then calculated and used to cluster the cells into k clusters. We used the function “kmeans” from the base R package “stats”.

### Cluster validation with R

Ideal clusters will be compact, well separated and connected. In other words, we want to minimize the intra-cluster variation, maximize the inter-cluster distances, and that each object and its nearest neighbors are in the same clusters [55]. Compactness tends to increase with cluster size, whereas separation and connectedness decrease. There are many metrics that combine them and can be used to quantitively compare different clustering methods and to determine the ideal number of clusters.

To validate our clustering results, we used 6 different metrics: BW ratio, Dunn index, average Silhouette width, Pearson correlation index, Calinski and Harabasz index (or variance ratio), and Connectivity. We define BW as the ratio between the average of all distances between elements of different clusters, and the weighted average (to cluster size) of averages of distances between elements within a cluster. The connectivity, using a neighborhood size of 25, was calculated using the function “connectivity” from the package “clValid” [56]. The rest of metrics were calculated using the function “cluster.stats” from the package “fpc” [57].

### Data visualization

The data was visualized with R and the ggplot2 package. The scripts to produce the figures in the main text are available: https://github.com/JoachimGoedhart/GPCR-KTR

## Data & Code availability

The following plasmids are deposited at addgene (http://www.addgene.org/): #129631: H2A-mScarletI-P2A-Akt-KTR-mTurquoise2-P2A-ERK-KTR-mNeonGreen, #129632: PB-H2A-mScarlet-2A-AKT_KTR-mTq2-2A-ERK_KTR-mNG Scripts for the analysis are available on GitHub: https://github.com/JoachimGoedhart/Nuclear-translocation-analysis Fully reproducible figures are available with the scripts for data visualization: https://github.com/JoachimGoedhart/GPCR-KTR

The data is present in the GitHub repo when file size limit allows. All source data will be deposited at zenodo.org upon acceptance of the manuscript.

## Author contribution

S.C.A. designed, analyzed and performed experiments and wrote the manuscript; M.L.B.G. generated constructs and performed microscopy experiments; T.W.J.G. & F.J.B. provided valuable ideas and input; J.G. designed experiments, assisted with the analysis and co-wrote the manuscript.

## Acknowledgements

We thank Ronald Breedijk for the support at the van Leeuwenhoek Centre for Advanced Microscopy, Section Molecular Cytology, Swammerdam Institute for Life Sciences, University of Amsterdam. We are grateful to Katrin Wiese and Anna Chertkova for assistance with FACS sorting. We thank Kazuhiro Aoki for sharing the pHGEA plasmid.

## Supplemental notes

Supplemental note 1 – Characterization of monoclonal cell lines

To test the dynamic range of the KTRs in the sorted cells, we used 5% FBS, given the strong stimulatory effect of the growth factors it contains on ERK and Akt activities. As it has been reported that the dynamic range of the KTRs can be negatively correlated to expression levels [10], we measured the FBS responses of several pools. We found no correlation between expression level (inferred from fluorescence intensity) and response but observed that some of the brightest cells displayed lower responses (data not shown). Therefore, we continued with cells from a pool with intermediate brightness.

For each of 13 monoclonal lines derived from this pool, we quantified the cellular fluorescence intensity prior to stimulation, and the translocation of the Akt- and ERK-KTRs in response to serum. The results, in Supplemental Figure S3, show the heterogeneity of the responses, both between single cells within a monoclonal population, and between clones. Overall, the maximum change in the C/N ratios are significantly higher for the ERK-KTR than for the Akt-KTR. The average change for ERK is 0.53, which is 70% higher than the average change of 0.31 for Akt. Again, we do not observe any correlation between brightness and dynamic range (Supplemental figure S3C). We selected 5 clones for further characterization and examined their response to G protein-coupled receptors. Stimulation with high concentrations of histamine (100 μM), S1P (1.3 mM), and UK (10 nM), showed clear translocation of both KTRs in all 5 clones, except for clone D3 which did not respond to UK. We decided to use clone E2 for further studies with these ligands due to higher brightness than the other clones.

### Supplemental tables

**Supplemental Table T1.** Brightness and dynamic range of the 13 monoclonal populations originated from pool 3. The C/N change for Akt- and ERK-KTRs in response to 5% FBS is the average of the maximum C/N change of all the cells per clone across the biological replicates. Per cell, the maximum C/N change is the highest C/N ratio after stimulation with 5% FBS, normalized by subtracting the average C/N ratio prior to stimulation. Brightness in the mTq2 and mNG channels is expressed as the average of average cellular fluorescence from various biological replicates. For each replicate, the average was normalized to the average of two replicates of pool 3 in the same slide. For each cell and channel, the cellular fluorescence intensity was calculated as the average between time points 1 and 7, prior to stimulation with 5% FBS. n: number of biological replicates. In bold and blue, the 5 clones selected for further characterization.

| Clone | # cells | C/N ratio change |  | Brightness |  | n |
| --- | --- | --- | --- | --- | --- | --- |
|  |  | Akt-KTR | ERK-KTR | mTq2 | mNG |  |
| A6 | 2327 | 0.4 | 0.4 | 1.4 | 1.3 | 3 |
| **B3** | **2008** | **0.31** | **0.62** | **0.86** | **0.85** | **5** |
| B5 | 1267 | 0.28 | 0.67 | 0.65 | 0.63 | 4 |
| C4 | 2054 | 0.24 | 0.6 | 1.14 | 1.42 | 5 |
| **D3** | **2346** | **0.32** | **0.57** | **1.55** | **1.92** | **3** |
| D8 | 812 | 0.22 | 0.41 | 0.94 | 0.83 | 3 |
| **E2** | **2138** | **0.3** | **0.62** | **0.91** | **0.84** | **3** |
| **F6** | **1679** | **0.32** | **0.59** | **0.89** | **0.93** | **3** |
| F11 | 1709 | 0.29 | 0.52 | 0.78 | 0.73 | 3 |
| F12 | 1851 | 0.28 | 0.51 | 0.94 | 0.89 | 3 |
| G2 | 1196 | 0.22 | 0.37 | 1.1 | 1.04 | 3 |
| G7 | 1192 | 0.3 | 0.41 | 1.19 | 1.18 | 4 |
| **G10** | **1150** | **0.33** | **0.5** | **1.03** | **1.14** | **4** |
| Pool 3 | 17466 | 0.35 | 0.48 | 1.0 | 1.0 | 16 |

**Supplemental Table T2.** Parameters that describe the concentration-response curves fitted in Figure 3 using ERK AUC as the measure of response. Half maximal effective concentration (EC50), Hill slope, and upper and lower limits.

| Condition | Ligand | EC50 | Hill | Lower | Upper |
| --- | --- | --- | --- | --- | --- |
| DMSO | His | 0.28 μM | 0.63 | -0.60 | 2.97 |
| PTx | His | 0.41 μM | 0.68 | -0.43 | 3.30 |
| YM | His | 0.03 μM | 0.52 | -0.78 | 0.87 |
| YM+PTx | His | 0.03 μM | 1.99 | -0.33 | 0.66 |
| DMSO | S1P | 64 nM | 1.49 | -0.12 | 5.02 |
| PTx | S1P | 61 nM | 0.66 | -0.50 | 2.84 |
| YM | S1P | 140 nM | 0.95 | -0.16 | 6.02 |
| YM+PTx | S1P | 36 nM | 0.48 | -1.33 | 3.73 |
| DMSO | UK | 2.1 pM | 0.89 | -0.11 | 2.99 |
| PTx | UK | 0.34 pM | 7.89 | -0.03 | 0.24 |

**Supplemental Table T3.** Sum of validation metrics for all the candidate clustering methods. Per kinase, the values from each of the six metrics were normalized by dividing them by the maximum score among the 15 combinations, and the sum of the six normalized metrics is shown. In bold and blue, the two highest scores per kinase.

| Method | Akt |  | ERK |  |
| --- | --- | --- | --- | --- |
|  | k | Sum | k | Sum |
| **eucl_kmns** | **3** | **5.46** | 8 | 5.35 |
| eucl_kmns | 4 | 5.27 | 9 | 5.18 |
| eucl_kmns | 5 | 4.93 | 10 | 4.88 |
| eucl_wrD1 | 3 | 4.78 | 8 | 4.94 |
| eucl_wrD1 | 4 | 5.00 | 9 | 4.26 |
| eucl_wrD1 | 5 | 4.05 | 10 | 4.02 |
| **eucl_wrD2** | **3** | **5.47** | **8** | **5.73** |
| eucl_wrD2 | 4 | 4.91 | 9 | 5.61 |
| eucl_wrD2 | 5 | 4.65 | 10 | 4.45 |
| manh_wrD1 | 3 | 4.66 | 8 | 4.26 |
| manh_wrD1 | 4 | 4.98 | 9 | 4.49 |
| manh_wrD1 | 5 | 4.82 | 10 | 4.42 |
| **manh_wrD2** | **3** | **5.11** | **8** | **5.53** |
| manh_wrD2 | 4 | 5.24 | 9 | 5.44 |
| manh_wrD2 | 5 | 4.70 | 10 | 4.61 |

### Supplemental figures

**Supplemental Figure S1.**
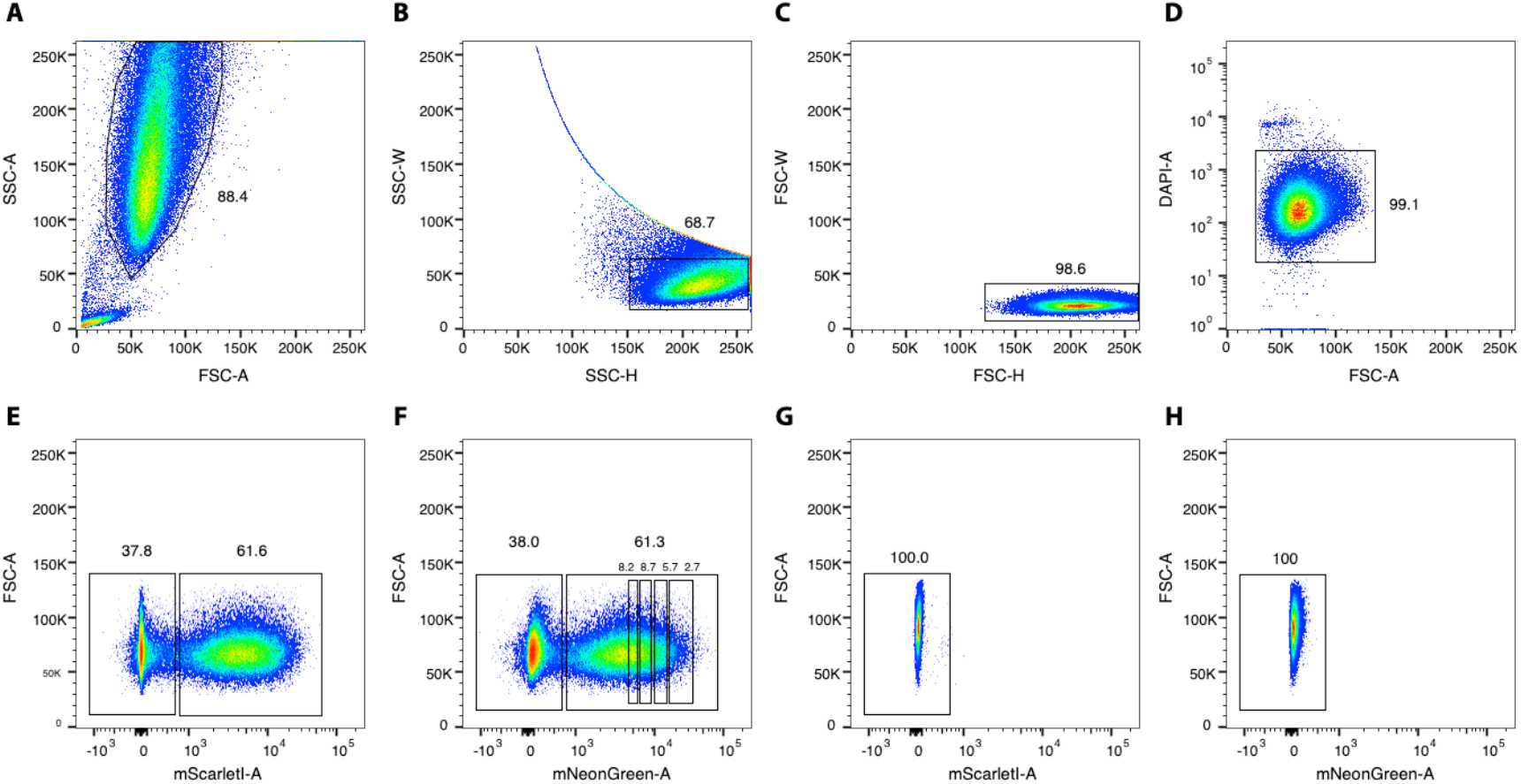
FACS gating for HSATEN cells. A-F: HeLa cells stably transfected with PB-HSATEN post puromycin selection. A-C: Gating for single cells based on forward (FSC) and side scatter (SSC). D: Gating for living cells based on DAPI staining. E: Gating for mScI positive cells. F: Gating for mNeonGreen positive cells. G-H: HeLa controls cells. G: Gating for mScI negative cells. H: Gating for mNeonGreen negative cells.

**Supplemental Figure S2.**
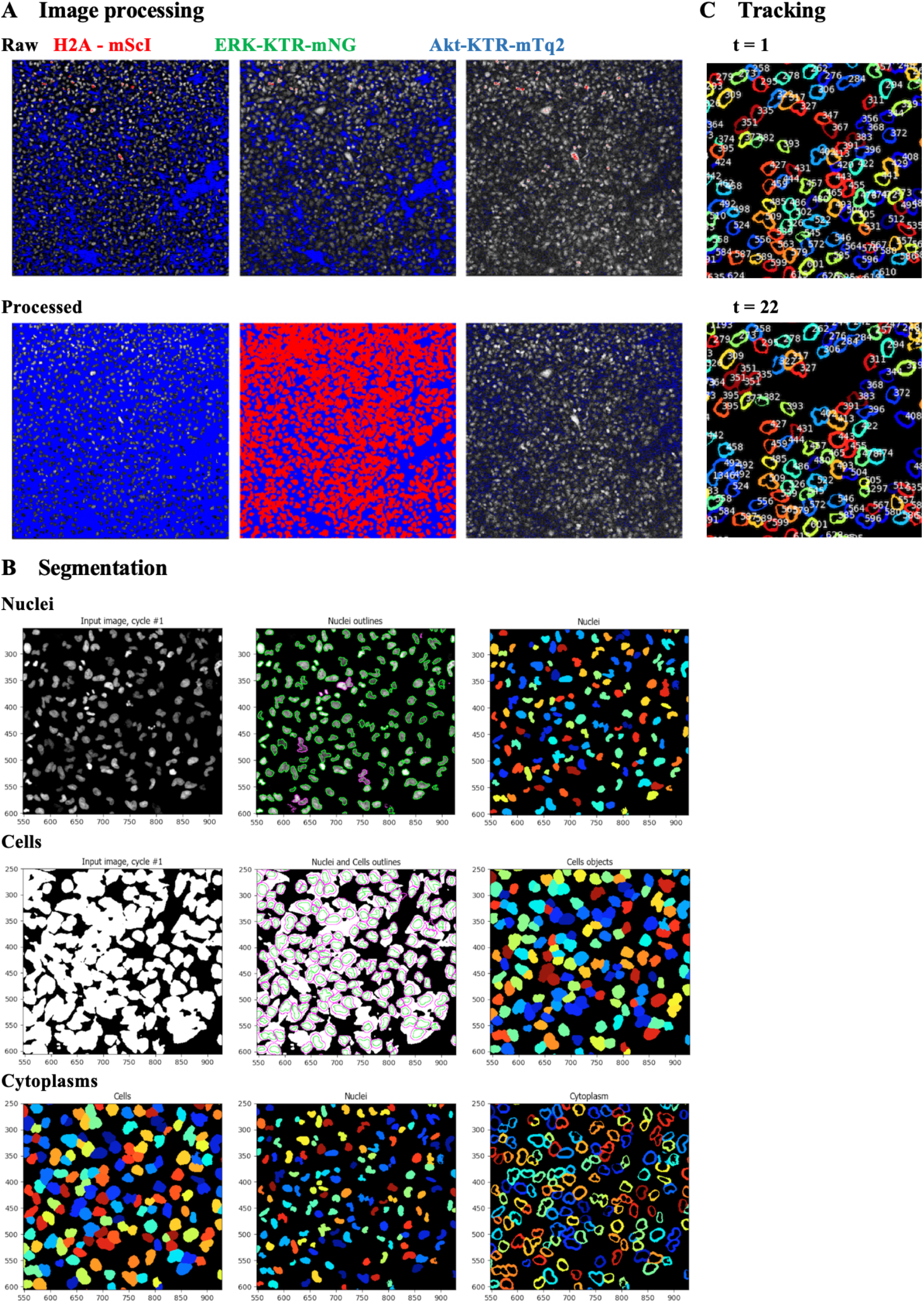
Image processing, segmentation and tracking. A: Background subtraction and noise reduction of raw images with FIJI. Top: Raw images of the mScI, mNG and mTq2 channels from a monoclonal population of HeLa cells stably expressing HSATEN. Bottom: Top images after processing. For mScI, we subtracted 250 counts. For mNG, we applied a Gaussian blur with sigma 2 and a threshold from 300 to 65535 to create a binary mask. For mTq2, we used the ‘Subtract Background’ function in FIJI with a rolling ball of 70 pixels. All panels are visualized using a HiLo LUT, which displays the dimmest pixels as blue, the brightest pixels as red, and the rest as a grayscale. B: Segmentation of nuclei, cells, and cytoplasms in CellProfiler. The panels show zoomed areas of the entire field of view. For the nuclei: Left panel shows the nuclear input image, which is the processed mScI image from panel. Center panel shows segmented nuclear outlines, with accepted objects in green and discarded objects in purple. Right panel shows segmented nuclei. For the cells: Cellular segmentation uses the segmented nuclei as seeds. Left panel shows the input image, which is the binary mask from in panel. Center panel shows segmented cellular outlines, with nuclear objects in green and cellular areas in purple. Right panel shows segmented cells. For the cytoplasms: Cytoplasmic segmentation results of subtracting the nuclear areas from the cellular areas. Left and center panels show the segmented nuclei and cells. Right panel shows segmented cytoplasms. C: Tracking of cytoplasms with CellProfiler. The left panels show the tracking of cells in a single field of view at the first and last time points of a time lapse of 22 images. The right panels are zoomed regions. The nucleus and cytoplasm of a single cell is identified with a unique tracked object number, shown next to each cytoplasm outline.

**Supplemental Figure S3.**
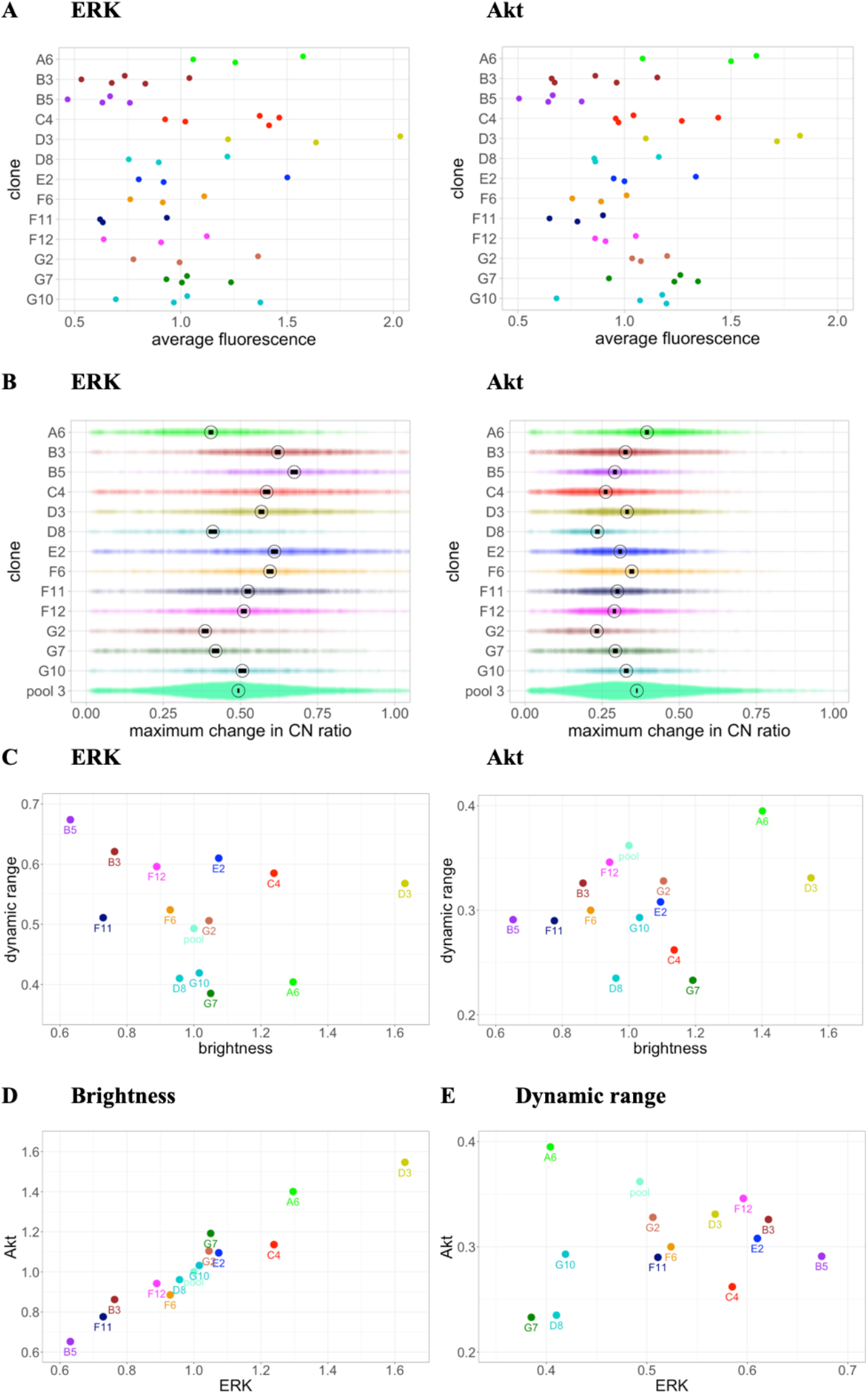
Comparison of brightness and dynamic range of the 13 monoclonal populations originating from pool 3. A: Cellular fluorescence (arbitrary units) in the mTq2 (Akt-KTR) and mNG (ERK-KTR) channels. Each dot represents the average cellular fluorescence intensity in a biological replicate. For each cell and channel, the cellular fluorescence intensity was calculated as the average between time points 1 and 7, prior to stimulation with 5% FBS. B: Maximum change in C/N ratio for the Akt- and ERK-KTRs in response to 5% FBS. Each dot represents a single-cell value and corresponds to the highest C/N ratio after stimulation with 5% FBS. Each value is normalized by subtracting the average C/N ratio prior to stimulation. For each clone, the mark and the circle represent the mean and 95% CI of the mean. Plots were generated using PlotsOfData [58]. C: The dynamic range of each clone plotted against the brightness for ERK and Akt respectively. D: Brightness of ERK and Akt per clone and E: the dynamic range of ERK-KTR versus that of Akt-KTR

**Supplemental Figure S4.**
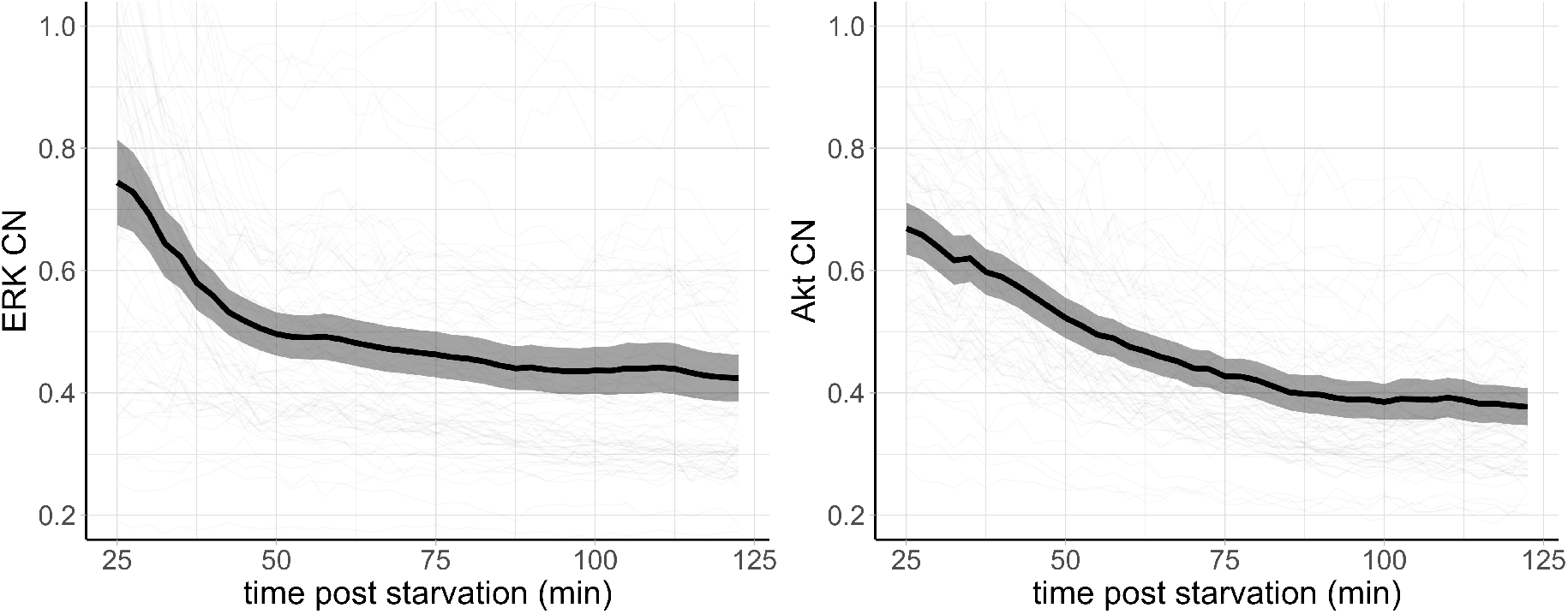
Effect of serum starvation on ERK and Akt C/N ratios. Each panel shows combined data from at least three biological replicates. The line shows the average and the ribbon shows the standard deviation.

**Supplemental Figure S5.**
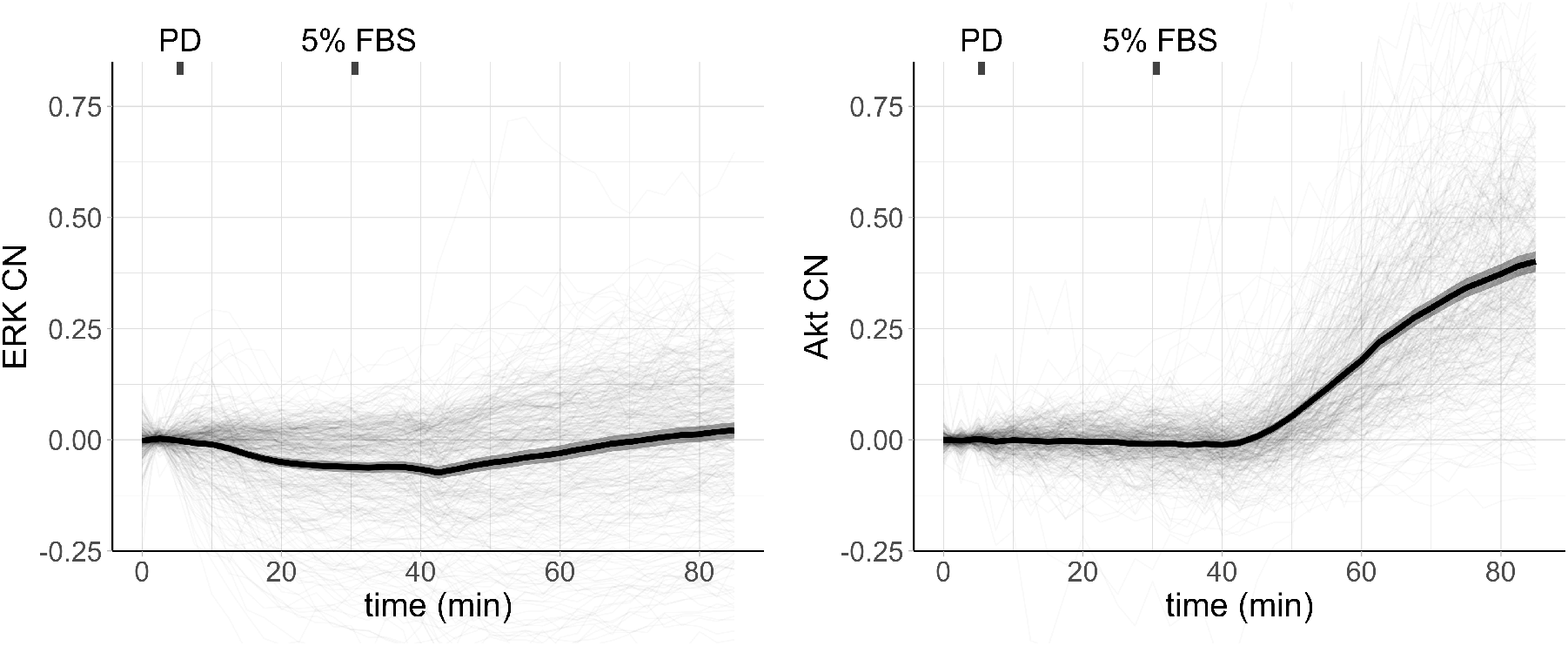
Effect of MEK inhibition on serum-dependent activation of ERK and Akt. 1 μM PD 0325901 and 5% serum were added at the indicated time-points. Each panel shows a representative experiment from at least three biological replicates. The line shows the average and the ribbon shows the standard deviation.

**Supplemental Figure S6.**
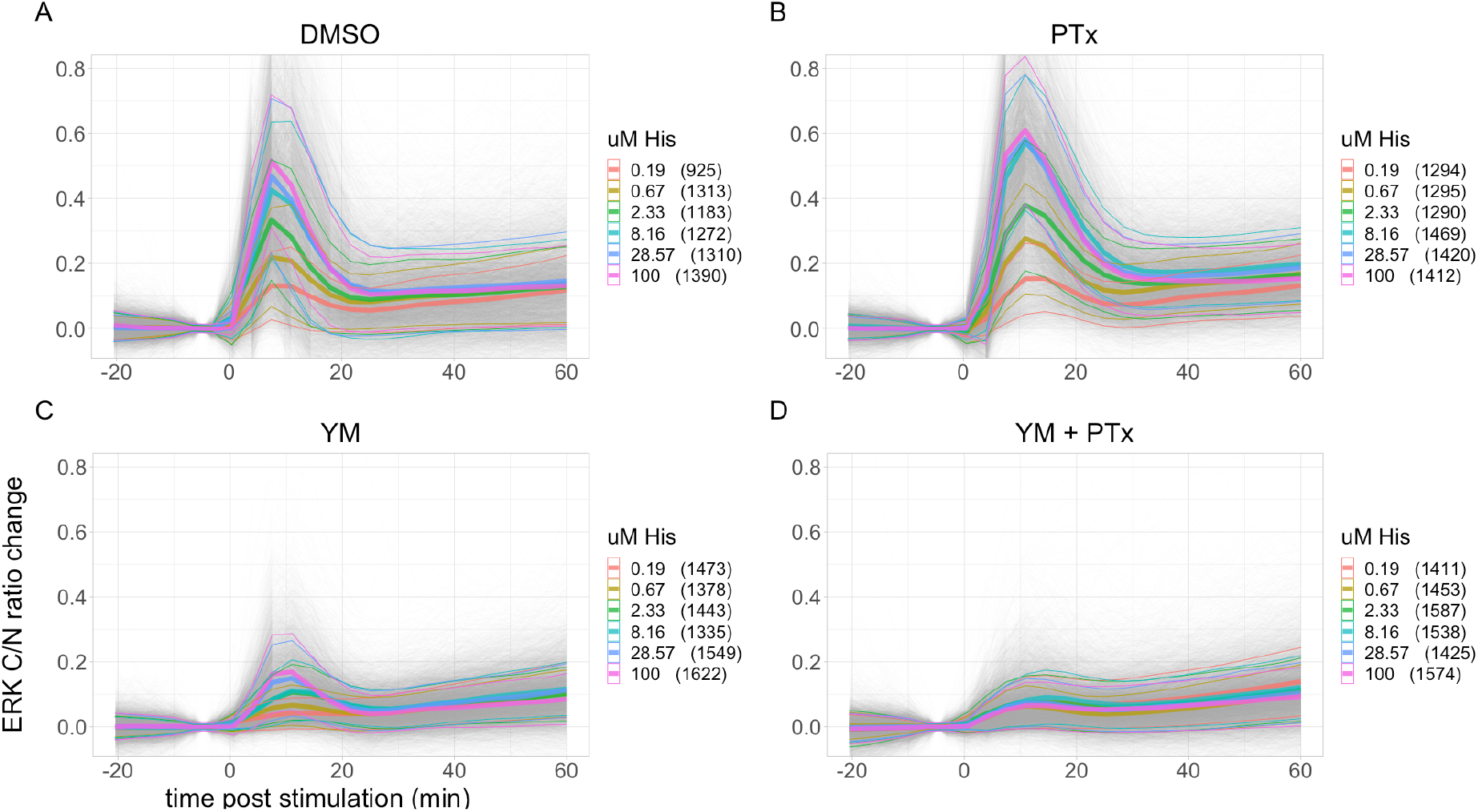
ERK responses to different concentrations of histamine and the effect of Gq and Gi inhibition. A: No inhibitor (DMSO). B: Gq inhibition (YM). C: Gi inhibition (PTx). D: Combined Gq and Gi inhibition (YM+PTx). ERK C/N ratio change is calculated by subtracting the average from the two time points prior to stimulation. Each panel shows combined data from at least three biological replicates. Gray lines represent single cell traces. Thick colored lines show the mean and thin colored lines the standard deviation for each ligand concentration. Number of cells are shown between brackets.

**Supplemental Figure S7.**
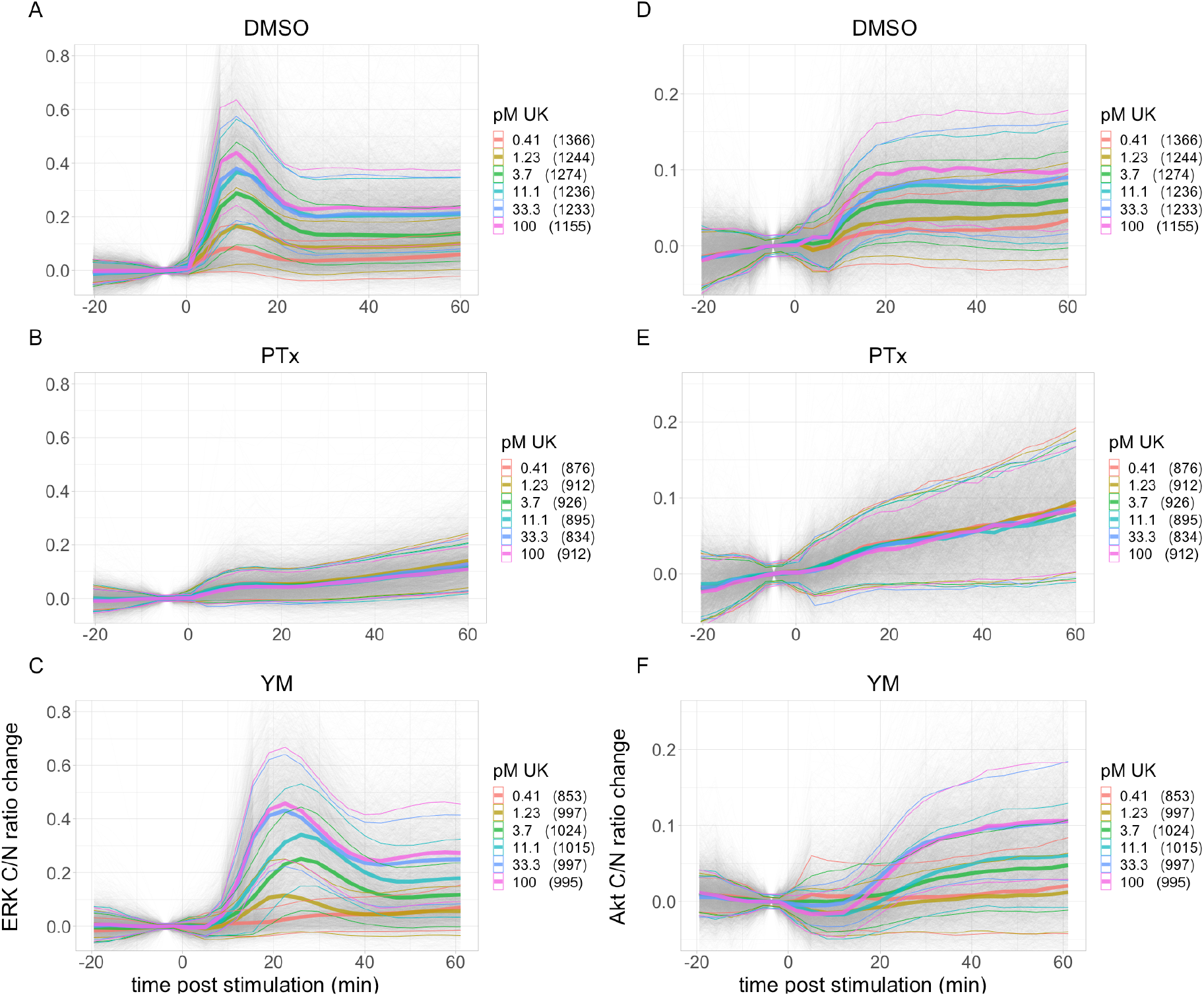
ERK and Akt responses to different concentrations of UK and the effect of Gi inhibition. A-B: ERK. C-D: Akt. A-C: No inhibitor (DMSO). B-D: Gi inhibition (PTx). ERK C/N ratio change is calculated by subtracting the average from the two time points prior to stimulation. Each panel shows combined data from at least three biological replicates. Gray lines represent single cell traces. Thick colored lines show the mean and thin colored lines the standard deviation for each ligand concentration. Number of cells are shown between brackets.

**Supplemental Figure S8.**
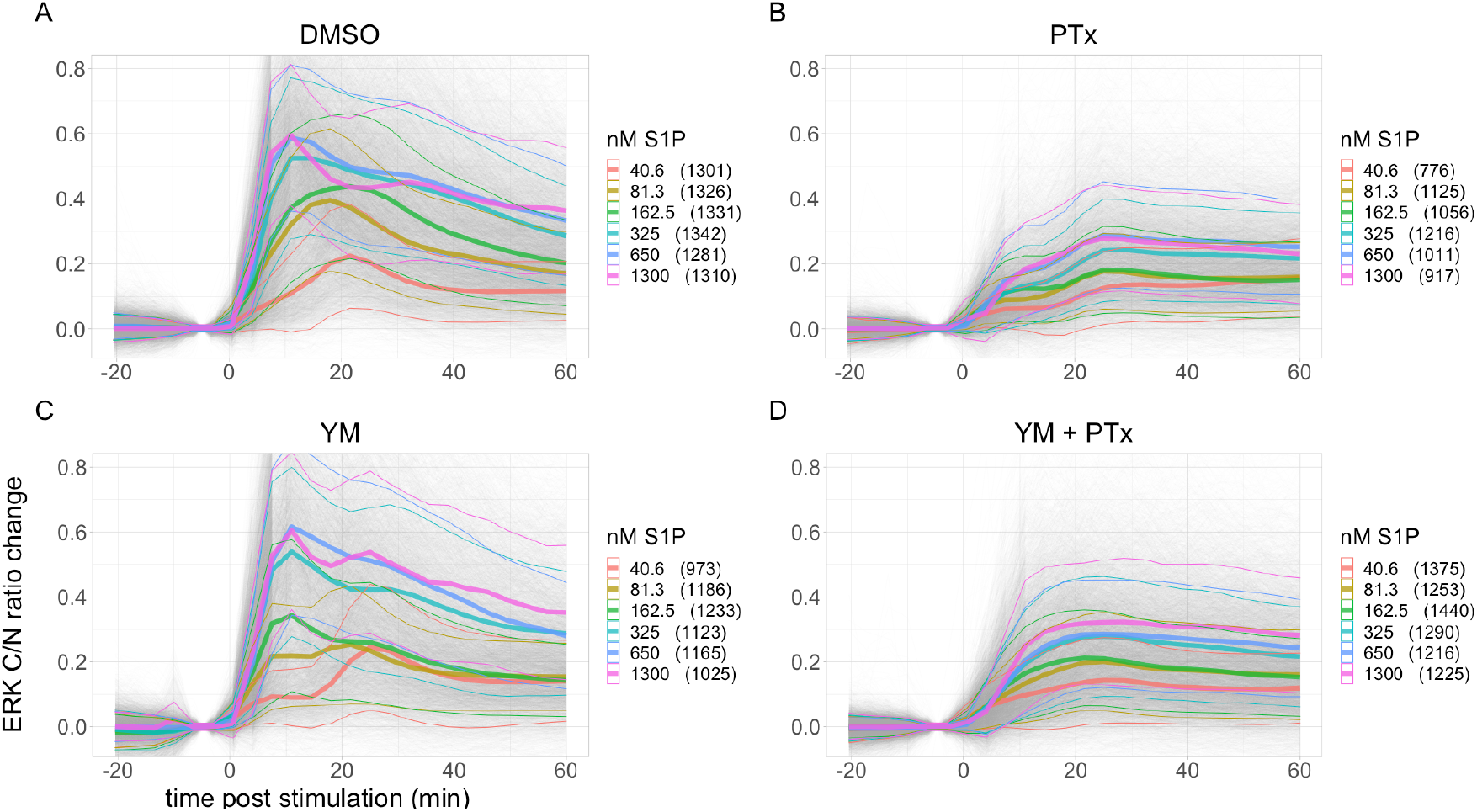
ERK responses to different concentrations of S1P and the effect of Gq and Gi inhibition. A: No inhibitor (DMSO). B: Gq inhibition (YM). C: Gi inhibition (PTx). D: Combined Gq and Gi inhibition (YM+PTx). ERK C/N ratio change is calculated by subtracting the average from the two time points prior to stimulation. Each panel shows combined data from at least three biological replicates. Gray lines represent single cell traces. Thick colored lines show the mean and thin colored lines the standard deviation for each ligand concentration. Number of cells are shown between brackets.

**Supplemental Figure S9.**
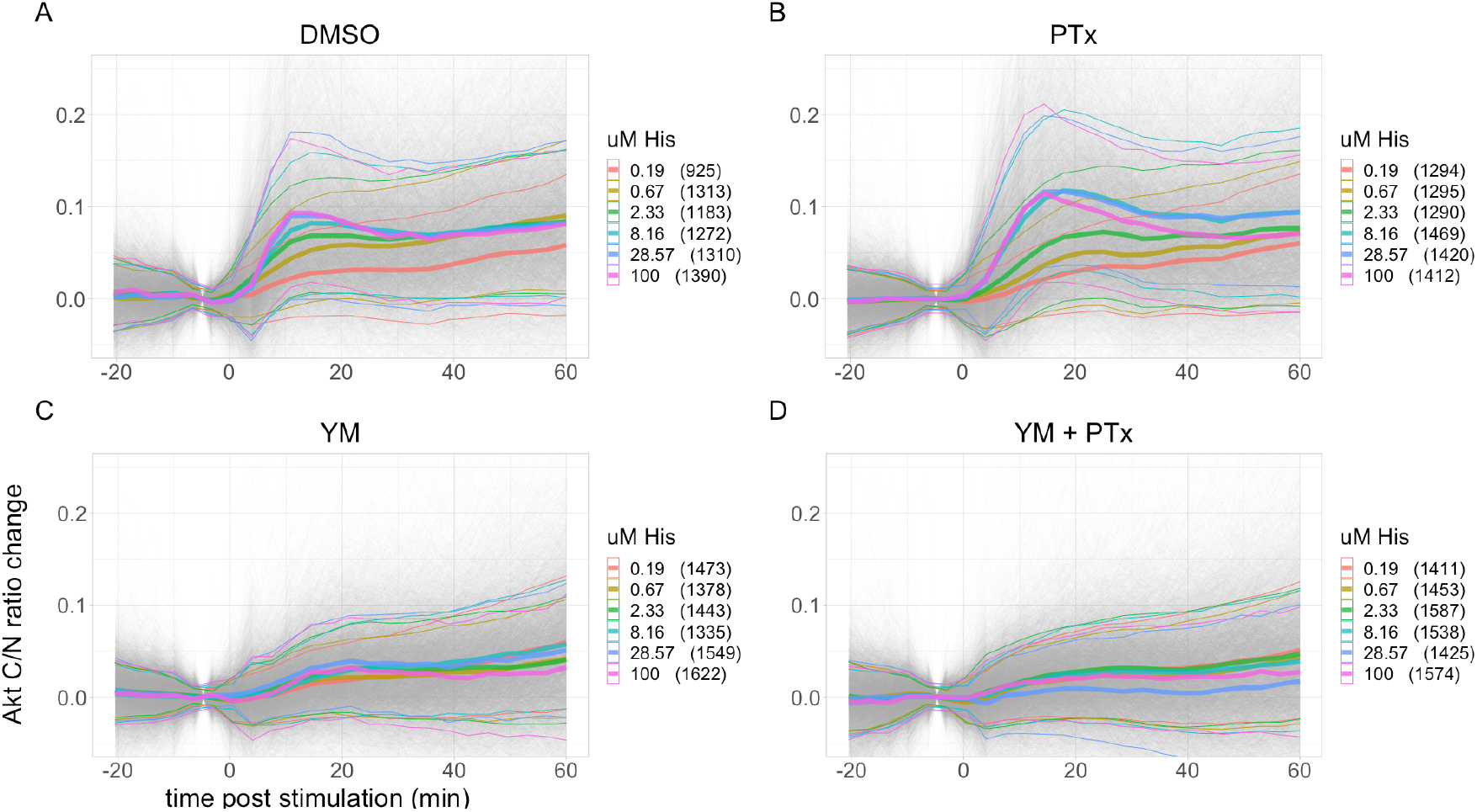
Akt responses to different concentrations of S1P and the effect of Gq and Gi inhibition. A: No inhibitor (DMSO). B: Gq inhibition (YM). C: Gi inhibition (PTx). D: Combined Gq and Gi inhibition (YM+PTx). ERK C/N ratio change is calculated by subtracting the average from the two time points prior to stimulation. Each panel shows combined data from at least three biological replicates. Gray lines represent single cell traces. Thick colored lines show the mean and thin colored lines the standard deviation for each ligand concentration. Number of cells per concentration are shown between brackets.

**Supplemental Figure S10.**
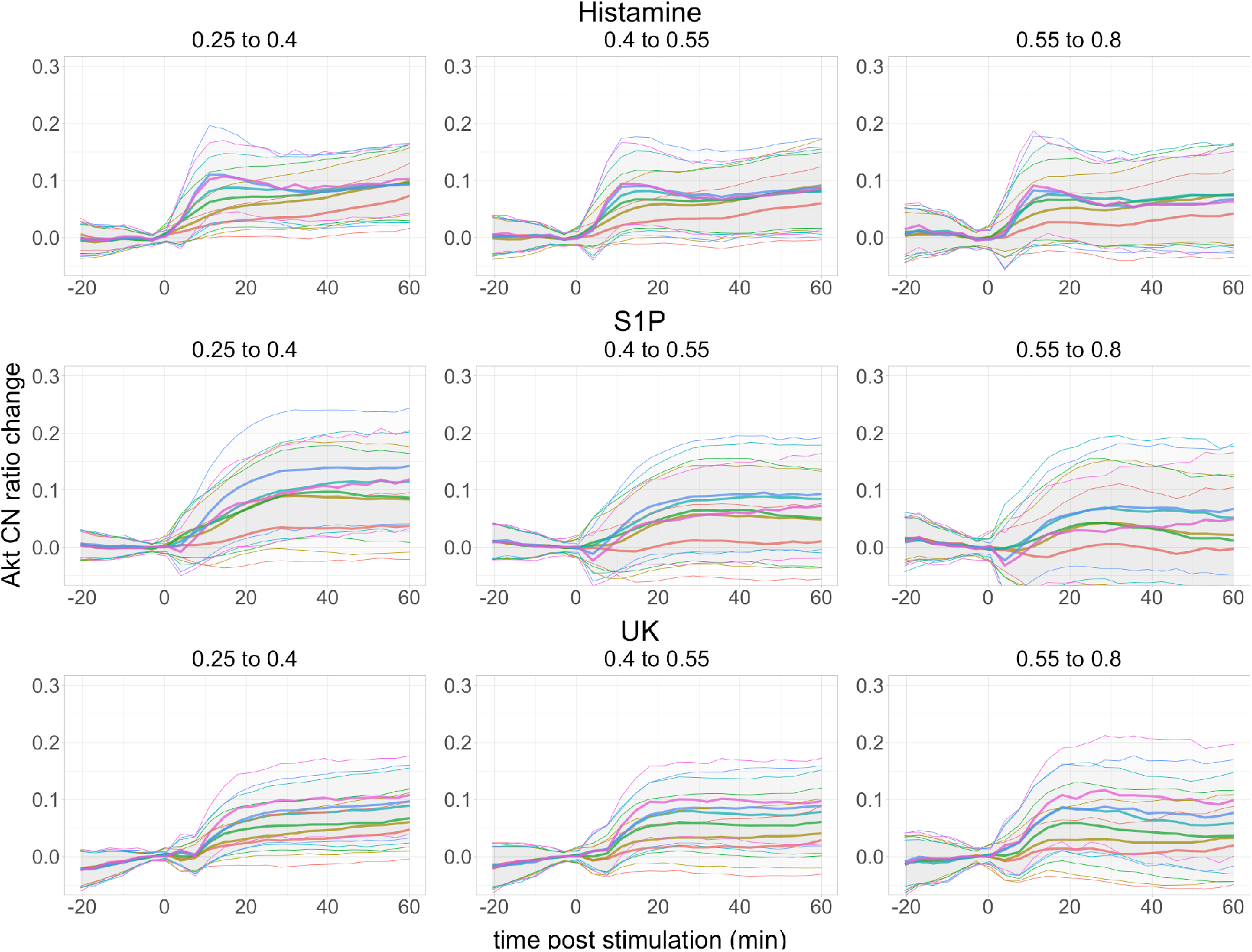
Effect of start ratio on average Akt responses upon multiple concentrations of histamine, UK, and S1P. Data from Figures 4A, S7C, and S9A, split in 3 groups according to the start-ratios, in the ranges 0.25-0.40, 0.40-0.55, and 0.55-0.80. The colors per ligand and concentration correspond to those shown in Figures 4, S7, and S9. The number of cells per concentration in the start-ratio groups shown from left to right, were in the following ranges: 174-260, 499-809, and 232-384 for histamine. 291-362, 630-780, and 156-216 for UK. 285-376, 750-799, and 189-245 for S1P.

**Supplemental Figure S11.**
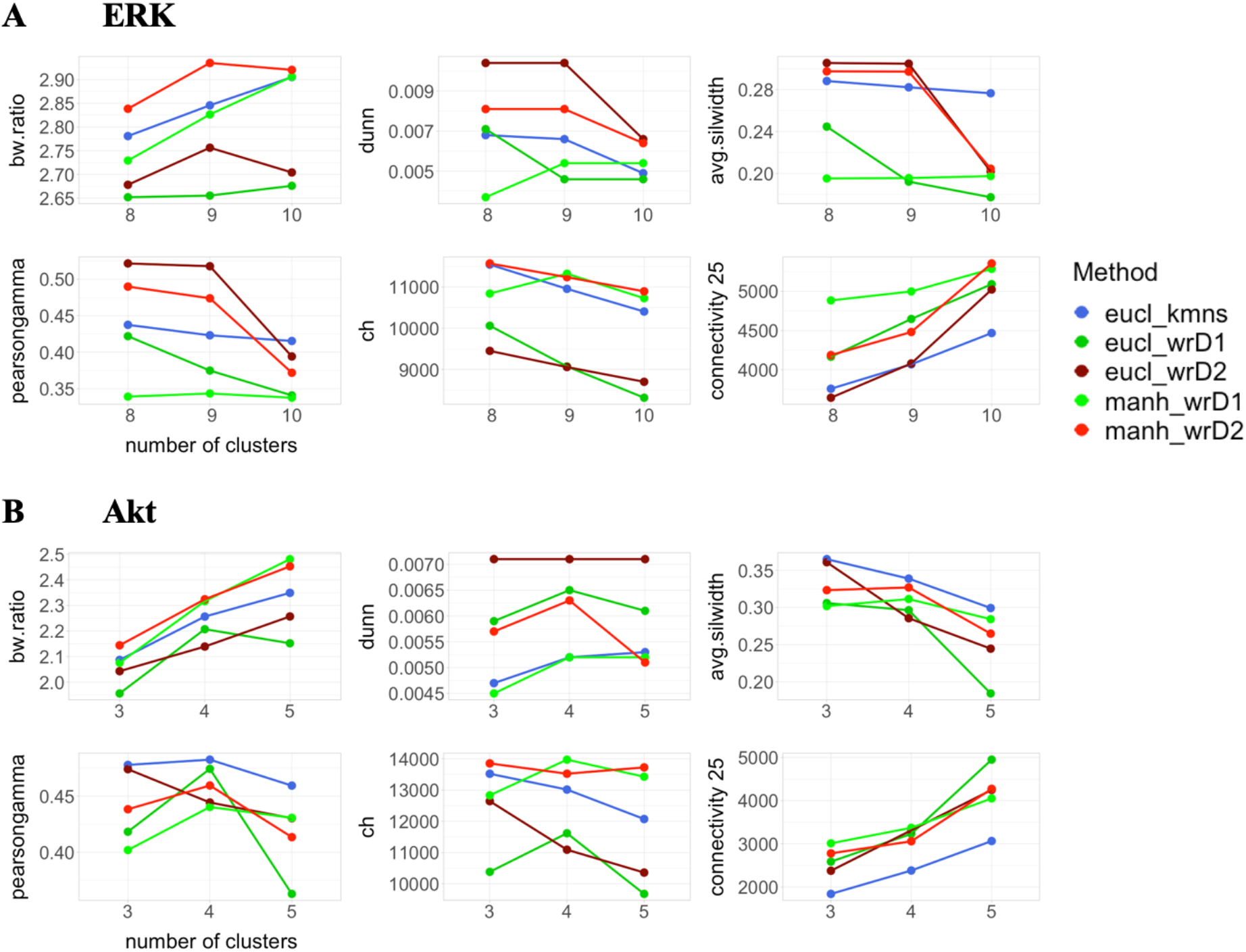
Validation metrics scores for all tested clustering methods with various number of clusters. A: For ERK, 8 to 10 clusters. B: For Akt, 3 to 5 clusters. Each clustering method was applied to a subset of 15 000 cells from the combined experiments with different ligands, concentrations, conditions, and negative controls. Negative controls include cells preincubated with 0.03% DMSO, 1 μM YM, or 100ng/mL PTx where microscopy medium was added instead of ligand. The validation metrics include the BW ratio, Dunn index, average Silhouette width, Pearson correlation index, Calinski and Harabasz index, and Connectivity. In blue: k-means clustering. In dark green: Euclidean distance and Ward linkage method. In dark red: Euclidean distance and Ward2 linkage method. In green: Manhattan distance and Ward linkage method. In red: Manhattan distance and Ward2 linkage method.

**Supplemental Figure S12.**
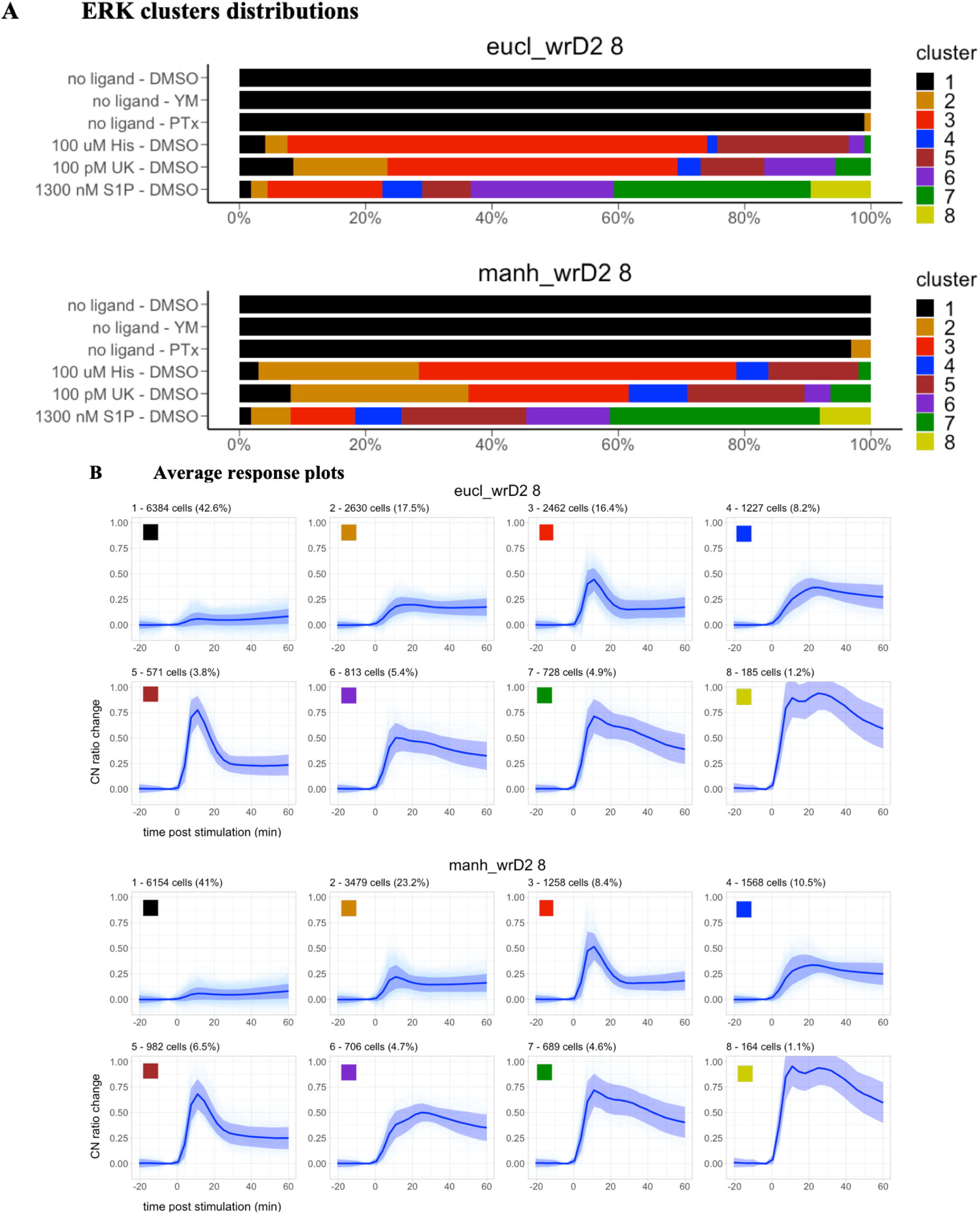
Clustering candidates for ERK responses. The two selected methods have 8 clusters, use Ward2 linkage method, and use the Euclidean or Manhattan distance. Each method was applied to a subset of 15 000 cells from the combined experiments with different ligands, concentrations, conditions, and negative controls. A: Cluster distribution of responses in negative and positive controls. Negative controls include cells preincubated with 0.03% DMSO, 1 μM YM, or 100ng/mL PTx where medium was added instead of ligand. Positive controls include cells preincubated with 0.03% DMSO where maximum stimulatory concentrations of Histamine, UK, and S1P were added. B: Average trajectory and frequency of each cluster. Per cluster, the lines represent the average trajectory and SD, and the number of cells and % of the total of 15 000 cells.

**Supplemental Figure S13.**
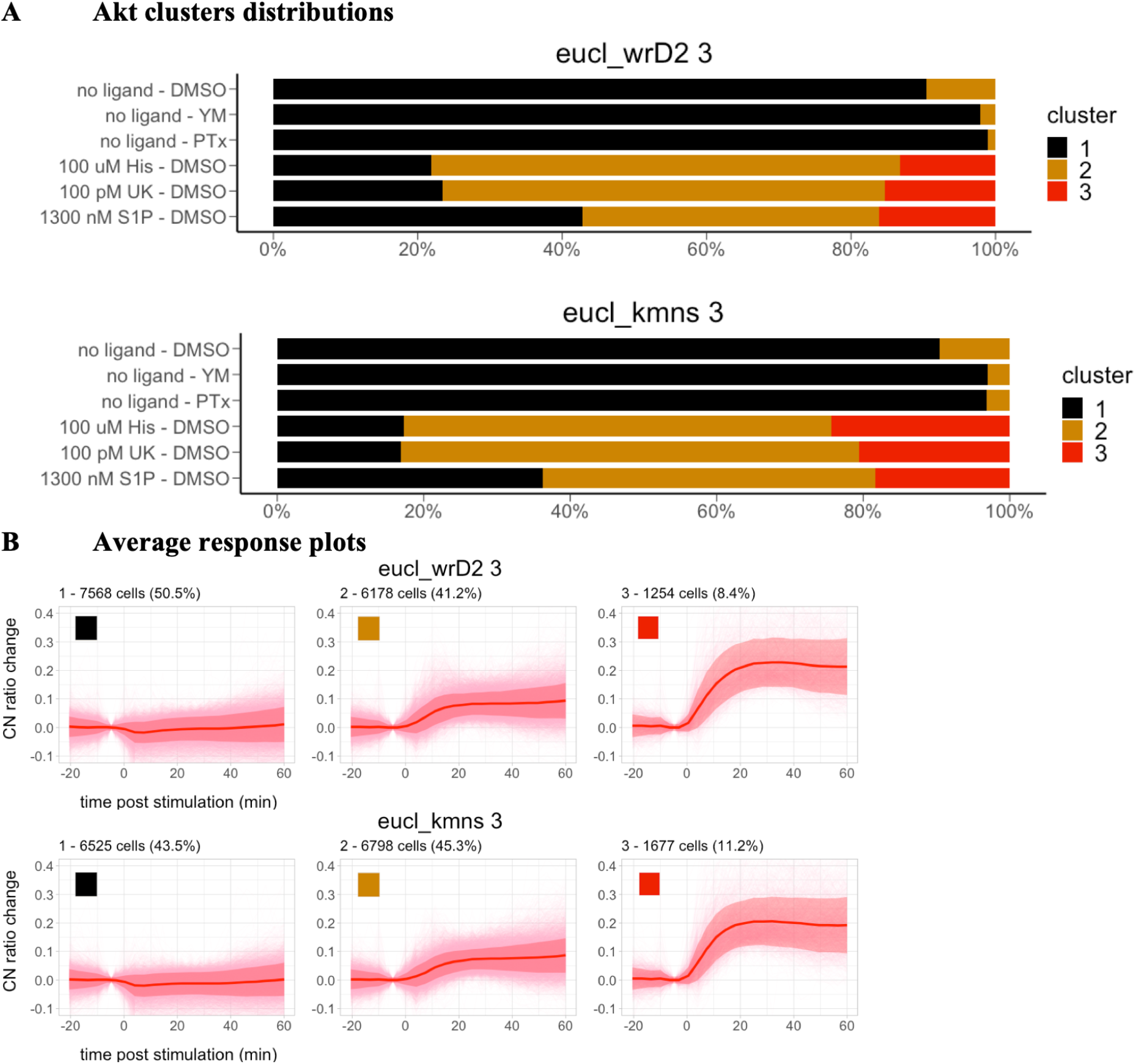
Clustering candidates for Akt responses. The two selected methods have 3-4 clusters, use Ward2 linkage method, and use the Euclidean or Manhattan distance. Each method was applied to a subset of 15 000 cells from the combined experiments with different ligands, concentrations, conditions, and negative controls. A: Cluster distribution of responses in negative and positive controls. Negative controls include cells preincubated with 0.03% DMSO, 1 μM YM, or 100ng/mL PTx where medium was added instead of ligand. Positive controls include cells preincubated with 0.03% DMSO where maximum stimulatory concentrations of Histamine, UK, and S1P were added. B: Average trajectory and frequency of each cluster. Per cluster, the lines represent the average trajectory and SD, and the number of cells and % of the total of 15 000 cells.

**Supplemental Figure S14.**
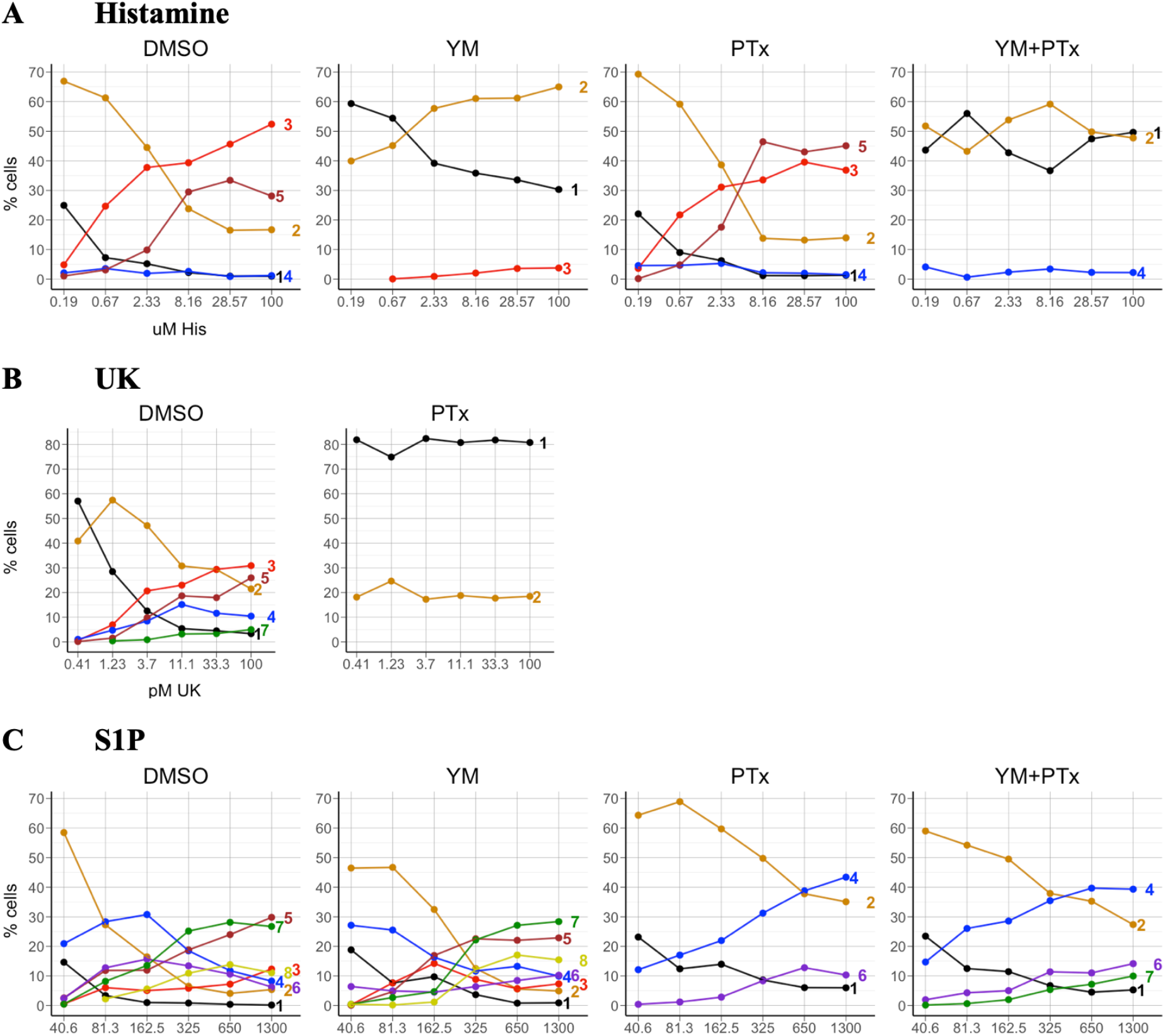
Cluster distribution of ERK responses per ligand in points. A. Histamine. B. UK. C. S1P. For each ligand, the panels represent the different experimental conditions. From top to bottom: No inhibitor (DMSO), Gq inhibition (YM), Gi inhibition (PTx), and combined Gq and Gi inhibition (YM+PTx). Only clusters with an average frequency of 2% or more across all concentrations are shown.

**Supplemental Figure S15.**
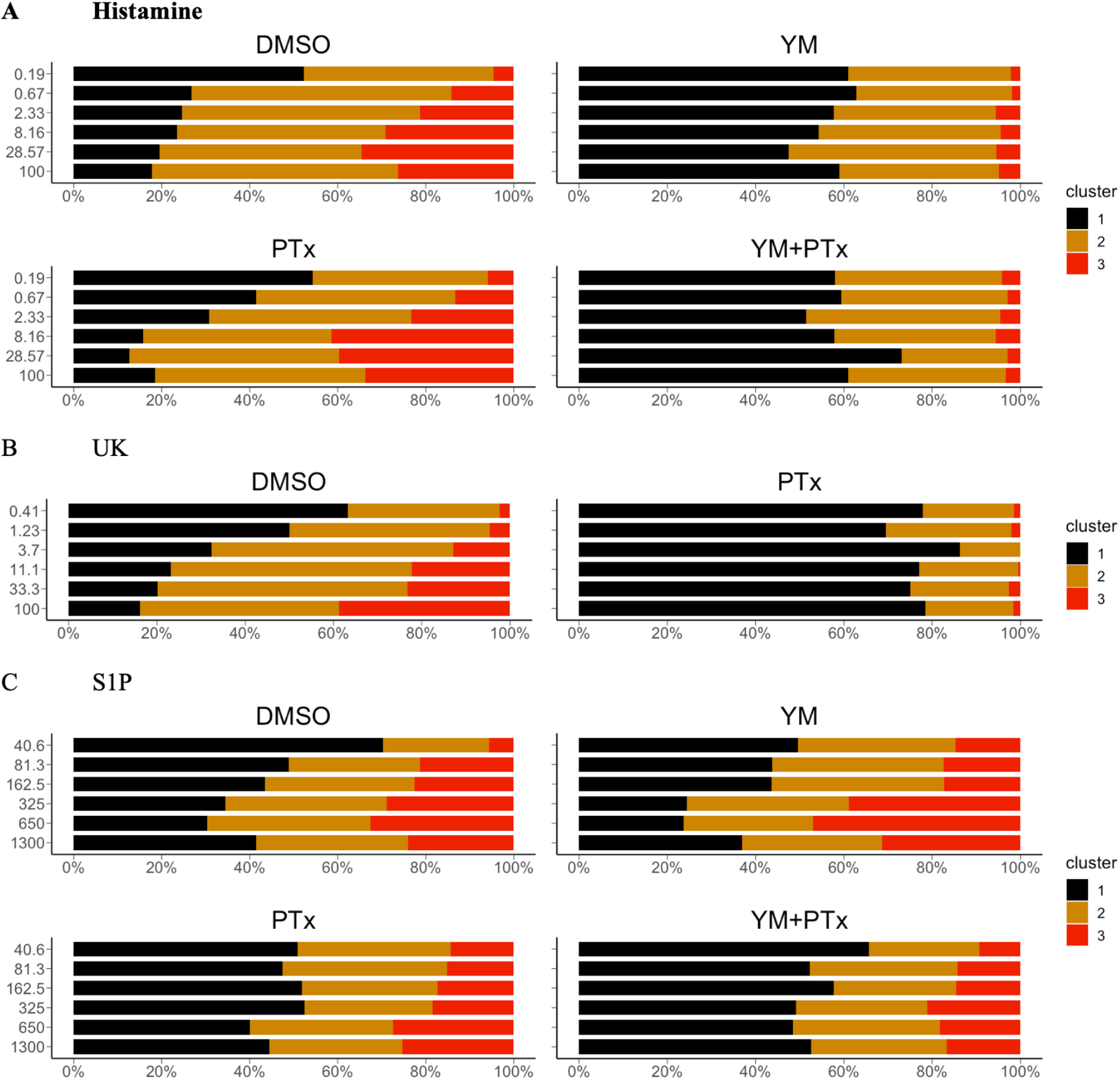
Cluster distribution of Akt responses per ligand in bars. A. Histamine. B. UK. C. S1P. For each ligand, the panels represent the different experimental conditions. From top to bottom: No inhibitor (DMSO), Gq inhibition (YM), Gi inhibition (PTx), and combined Gq and Gi inhibition (YM+PTx).

**Supplemental Figure S16.**
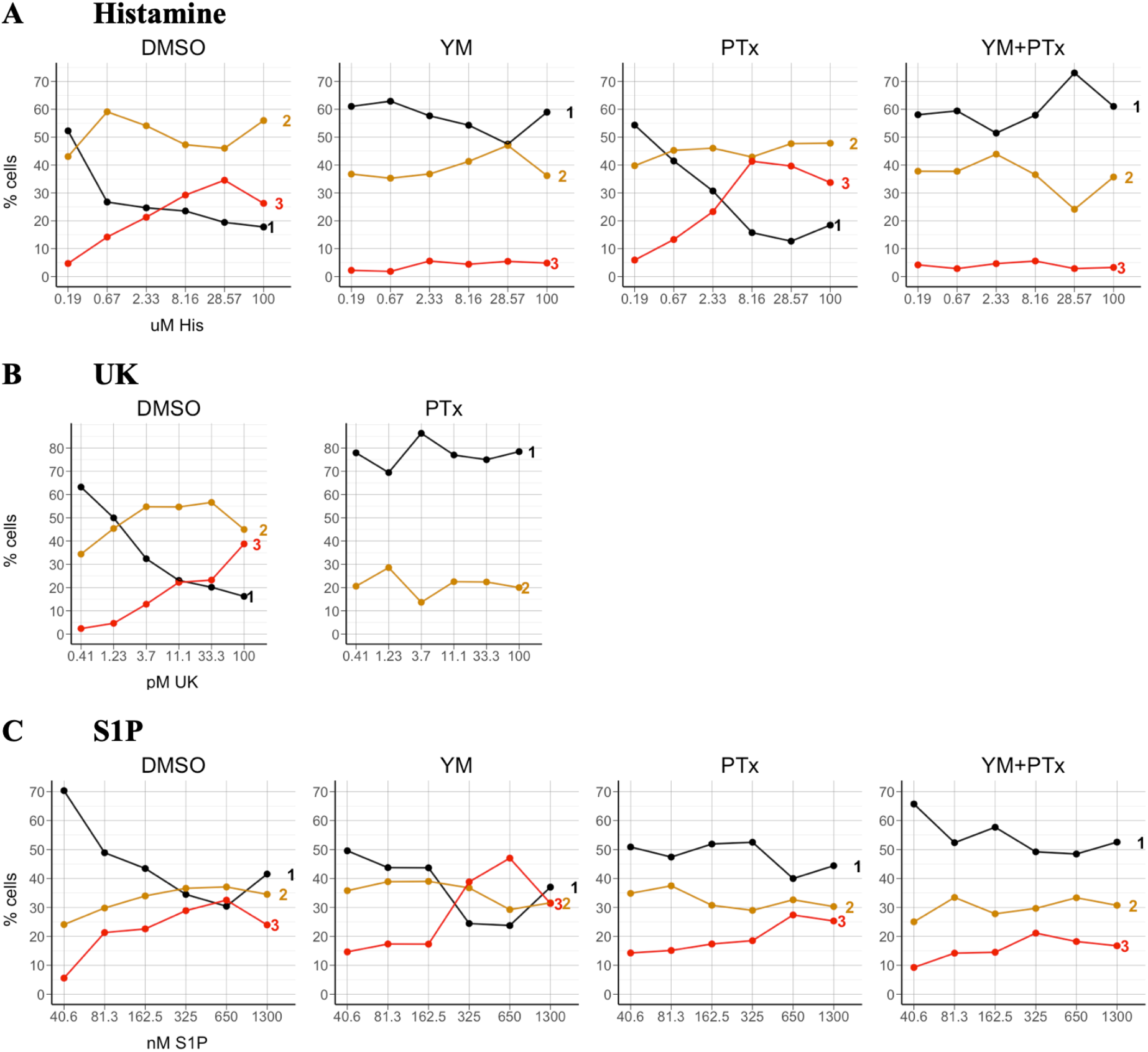
Cluster distribution of Akt responses per ligand in points. A. Histamine. B. UK. C. S1P. For each ligand, the panels represent the different experimental conditions. From top to bottom: No inhibitor (DMSO), Gq inhibition (YM), Gi inhibition (PTx), and combined Gq and Gi inhibition (YM+PTx). Only clusters with an average frequency of 2% or more across all concentrations are shown.

